# Prime-Target neoantigen vaccination unleashes unprecedented T cell immunity within “cold” immunosuppressive tumors

**DOI:** 10.64898/2026.01.13.699214

**Authors:** Kou Hioki, Marlous Wildemans, Youkyung Lim, Inge Brouwers–Haspels, Marjan van Meurs, Eric Bindels, Harmen J. G. van de Werken, Yvonne M. Mueller, Burcu Temizoz, Ken J. Ishii, Christopher Schliehe, Peter D. Katsikis

## Abstract

“Cold” immunosuppressive solid tumors are hard-to-treat cancers that are non-responsive to immunotherapies. Their immunosuppressive tumor microenvironment (TME) excludes and inhibits T cells and thereby hampers the therapeutic efficacy of cancer vaccines and immune checkpoint blockade. To overcome this, we employed a “Prime-Target” (P/T) neoantigen vaccination strategy that combines subcutaneous (SQ) and intra-tumor (IT) neopeptide vaccinations to first prime potent systemic neoantigen-specific T cell immunity and then trigger intratumoral T cell recruitment. Using immunotherapy non-responsive murine tumor models with pronounced immunosuppressive TMEs, we demonstrate that P/T neopeptide vaccination resulted in extraordinary tumor control and TME remodeling. P/T vaccination elicited strong systemic anti-tumor responses as well as potent and rapid recruitment of clonal CD4^+^ Th1 and non-exhausted CD8^+^ T cells into tumors that carried novel T cell receptors (TCR) and were vaccine neopeptide-specific. Concurrently, P/T vaccination reshaped the TME by decreasing suppressive Treg and M2 macrophages, and dramatically increasing the ratio of effector T cells to Treg and M2 macrophages. Vaccination-induced tumor control was neopeptide-dependent and required concurrent IT administration of both neopeptides and adjuvants. Our study highlights P/T neoantigen vaccination as a promising strategy to overcome T cell exclusion within “cold” immunosuppressive solid tumors and thereby unleash unprecedented anti-tumor immunity.

## INTRODUCTION

Therapeutic cancer vaccines hold great promise, but have yet to deliver clear benefit to cancer treatment^1^. One major strategy of therapeutic cancer vaccines targets tumor-specific neoantigens to boost pre- existing or induce *de novo* neoantigen-specific T cell responses systemically, and it employs different vaccine platforms such as mRNA-based and peptide-based vaccines^1^. However, these approaches until now are facing significant limitations, with only few patients responding favorably. One of the major obstacles to clinical success of cancer vaccines – and immunotherapies in general – is the immunosuppressive nature of the tumor microenvironment (TME) that is common across solid tumors^1^. This immunosuppressive TME interferes with anti-tumor immunity by impairing the intratumoral recruitment and function of T cells^2–4^. Therefore, merely boosting existing or inducing *de novo* anti-tumor T cell immunity on a systemic level is inadequate to effectively eliminate cancers. Thus, a successful therapeutic cancer vaccine must not only prime potent T cell immunity, but is additionally required to promote the recruitment of neoantigen-specific T cells into the TME and preserve their functional activity to ensure efficient cancer cell elimination^1^.

“Cold” solid tumors are resistant to immunotherapy and this is due to their lack of inflammation and sparse T cell infiltration^3,4^. While CD8⁺ T cells, which directly kill tumor cells in an antigen-specific manner, have traditionally been considered the primary effectors of cancer vaccine responses, increasing evidence indicates that CD4⁺ T helper 1 (Th1) cells also play crucial anti-tumor roles and associate with improved clinical outcomes of cancer immunotherapy ^5–7^. CD4^+^ Th1 cells can thereby orchestrate anti-tumor immune responses, including the effector function of CD8^+^ T cells, and re-shape the TME to a more favorable environment for T cell-mediated tumor elimination. Therefore, a successful cancer vaccine would most likely require to elicit both CD4^+^ and CD8^+^ T cell immunity, and convert a “cold” TME into a “hot” inflamed state with strong CD4^+^ Th1 and CD8^+^ T cell infiltration. This TME reshaping is a critical step toward enabling effective cancer vaccines and immunotherapy but remains an unresolved problem^1,8,9^.

To overcome the “cold” TME of cancers, a number of approaches have been taken. *In situ* vaccinations where immune stimulants/adjuvants without antigen are directly delivered into tumors have attempted to convert a “cold” to a “hot” TME, but shown little benefit in clinical trials^10,11^. Other attempts to directly alter the TME have employed dendritic cells (DC), oncolytic viruses or RNA^12^. Intranodal injections of DC or RNA directly into (draining) lymph nodes have also been employed to stimulate better T cell immunity^13–17^. These approaches, however, have failed to elicit sufficient protective immunity or TME activation in cancer patients. This gap highlights the need for therapeutic strategies capable of converting the TME into a proinflammatory state that supports tumor antigen-specific T cell recruitment and potent anti-cancer effector functions.

Independent of the challenges induced by the TME of “cold” tumors, an effective neoantigen vaccine needs to overcome the inferior immunogenicity of neoantigens and the lower affinity that often characterizes TCR that recognize these peptides that show sequence similarities with self-antigens^1^. We previously addressed this lower immunogenicity challenge by employing a potent dual-adjuvant combination, consisting of a Toll-like receptor 9 (TLR9) and a stimulator of interferon genes (STING) agonist. These adjuvants when combined with neoantigen peptides elicited 10-times higher neoantigen-specific T cells responses in mice, compared to the best-in-class T cell adjuvant, the TLR3 agonist poly I:C^18^. However, despite eliciting unprecedented numbers of neoantigen-specific T cells systemically in blood and spleens after subcutaneous (SQ) neoantigen vaccination^18^, we were unable to confer protection from “cold” tumors to animals, something we attributed to T cell exclusion from tumors and their immunosuppressive TME. Previously, we demonstrated that effector CD8^+^ T cells require antigen recognition, CD28 costimulation and dendritic cells (DCs) at the effector site for maximal expansion during viral infection^19^. The need for CD28 costimulation has also recently been confirmed for effector CD8^+^ T cells in the tumors^20^. These observations led us to combine our potent systemic SQ neopeptide vaccination with an intra-tumor (IT) vaccination of the same formulation to provide antigen and costimulatory/recruitment signals within the tumor, with the goal of circumventing T cell exclusion and achieving local intra-tumor T cell expansion and effector function. Based on its ability to first systemically prime neoantigen-specific T cells and then target these into the TME of “cold” tumors, we called this novel approach “Prime-Target” (P/T) vaccination.

To determine its efficacy, we evaluated our P/T neoantigen vaccination strategy in two immunotherapy non-responsive murine tumor models with an immunosuppressive TME phenotype: the AE17 mesothelioma model and the KPC-4662 pancreatic adenocarcinoma model. In these settings, the P/T neoantigen vaccination led to remarkably improved tumor control compared to conventional systemic SQ neoantigen vaccination, IT neoantigen vaccination alone or SQ neoantigen vaccination followed by IT adjuvants alone. This protection was accompanied by profound remodeling of intra-tumor T cell populations that after P/T vaccination were dominated by non-exhausted neoantigen-specific effector CD4^+^ (Th1) and CD8^+^ T-cells. Together, this work highlights the therapeutic potential of our P/T neoantigen vaccination strategy to greatly enhance neoantigen vaccine efficacy and unleash unprecedented anti-tumor immunity even in the setting of hard-to-treat “cold” solid tumors.

## RESULTS

### Prime-Target neoantigen vaccination controls immunosuppressive “cold” tumors

As mentioned above, we previously introduced a neopeptide vaccination adjuvanted with a combination of K3 CpG (a TLR9 agonist) and c-di-AMP (a STING agonist), referred to as K3/c-di-AMP^18^. Using subcutaneous (SQ) vaccination containing K3/c-di-AMP and 20 amino acid long neopeptides identified from mutations of murine AE17 mesothelioma, we could elicit strong systemic neopeptide-specific CD4^+^ and CD8^+^ T cell responses with *ex vivo* responses reaching average frequencies of 12.6% and 3.6% among CD4^+^ and CD8^+^ T cells in spleens, respectively (Extended Data Fig. 1a).

However, despite inducing potent systemic neopeptide-specific T cell responses, SQ vaccination alone did not control tumor growth in the AE17 tumor model (Extended Data Fig. 1b-d). This failure of tumor control by SQ vaccination was likely due to T cell exclusion and immunosuppression by AE17 tumors, as strong systemic neopeptide-specific T cell responses were present in these tumor-bearing animals (Extended Data Fig. 1e). Since we and others have shown that effector T cells require antigen and costimulation within effector sites and tumors^19,20^, we hypothesized that K3/c-di-AMP-adjuvanted IT neopeptide vaccination, which could promote tumor/TME-intrinsic neopeptide presentation and innate immune stimulation via TLR9 and STING, would potentiate the activity of SQ vaccine-induced T cells within the TME. As previously, we initially primed systemic T cell immunity in tumor-bearing mice by administering SQ neopeptide vaccinations. We then directed this immune response toward the tumors using IT neopeptide vaccination with the same formulation—a strategy referred to as Prime-Target (P/T) vaccination (Fig. 1a). Notably, we found that the mice receiving P/T neoantigen vaccination formed necrotic ulcers on their tumors indicating rapid tumor death (Fig. 1b). Both the frequency and size of ulcers were significantly greater than those found in mice that received either saline SQ and IT (SQ saline + IT saline, control) or SQ neopeptide vaccine in combination with IT K3/c-di-AMP adjuvants alone (SQ vaccine + IT adjuvants) (Fig. 1b). Necrotic ulcers began to appear as early as two to three days post 1^st^ IT neopeptide vaccination and have in general been positively correlated with increased efficacy of cancer immunotherapies^21–23^. In agreement with this, we observed that P/T neopeptide vaccination significantly suppressed tumor growth and improved survival of treated mice (Fig. 1c, d) with 40% of mice (6 out of 15) exhibiting complete tumor rejection (Fig. 1e). Of note, neither IT neopeptide vaccination alone (no SQ priming) nor IT administration of neopeptides alone (no adjuvants) or IT adjuvants alone (no neopeptides) after SQ neopeptide vaccination conferred protection (Fig. 1c-e), indicating that effective tumor control required both SQ and IT neopeptide vaccination in the AE17 tumor model. We further confirmed P/T neopeptide vaccination-induced tumor control in the “cold”, immunosuppressive KPC-4662 pancreatic adenocarcinoma tumor model (Fig. 1f-j). P/T neopeptide vaccination showed remarkable tumor control (Fig. 1g), while in long term experiments, 80% of mice achieved complete tumor rejection (Fig. 1j). As with the AE17 model, both neopeptides and adjuvants needed to be included in SQ and IT vaccinations for KPC-4662 tumor control (Fig. 1h-j). We next asked whether neoantigen-specific immunity was required for this protection. We confirmed that the P/T neopeptide vaccination strategy requires tumor-specific antigens to achieve tumor rejection, as P/T vaccination using irrelevant pathogen-derived peptides (both CD4 and CD8 epitopes) did not confer tumor control (Extended Data Fig. 2a). Taken together, our data demonstrate that the P/T neopeptide vaccination strategy results in unprecedented control of tumor growth in multiple hard-to-treat mouse tumor models.

**Fig. 1.**
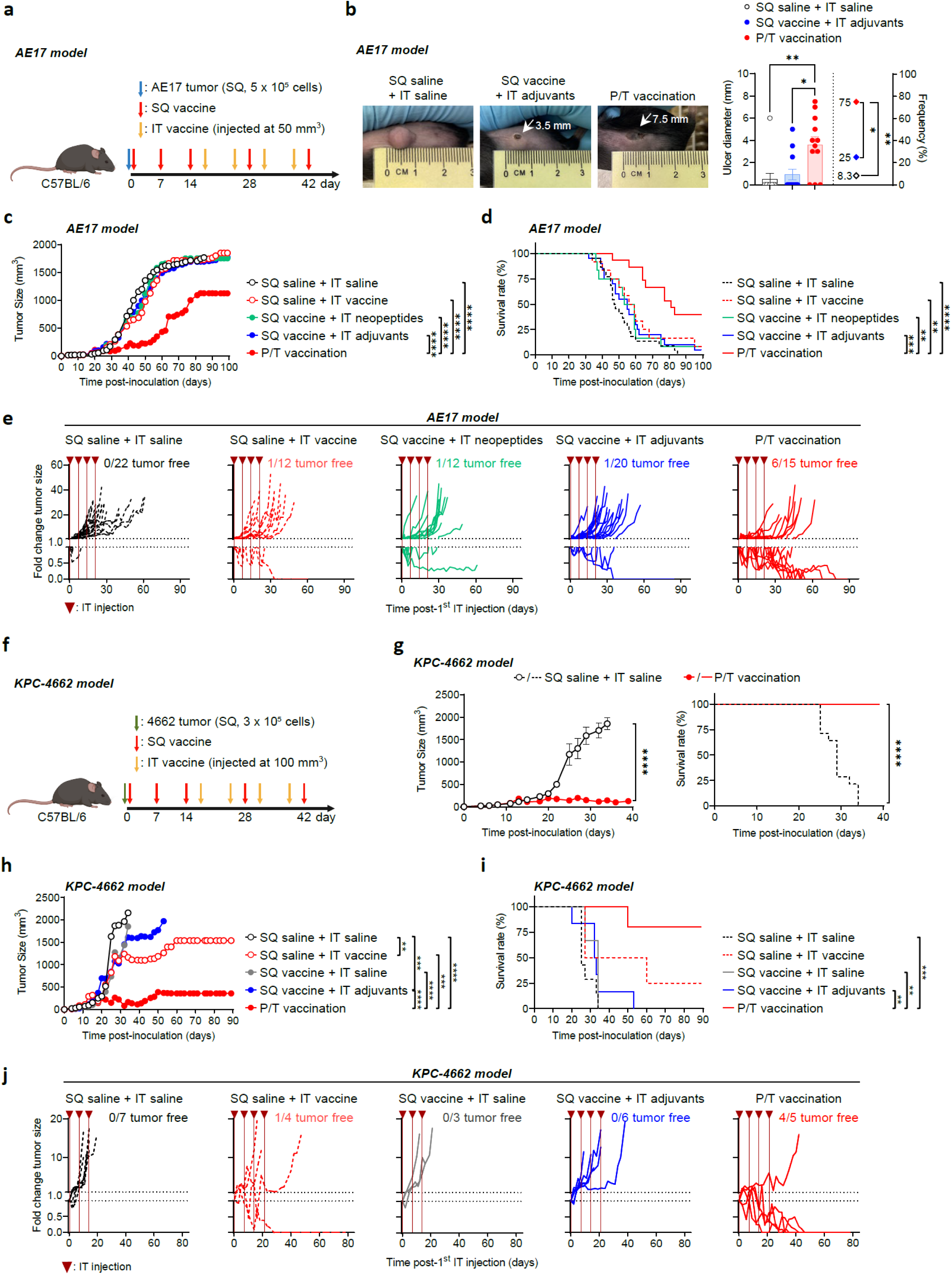
P/T neopeptide vaccination controls immunosuppressive tumors. **a**, Experimental design for P/T vaccination in the AE17 tumor model. C57BL/6J mice were inoculated with AE17 tumor cells (SQ, 0.5 x 10^6^ cells) on day 0. SQ neopeptide vaccines were administered on days 0, 7, 14, 28, and 42. Meanwhile, tumor-bearing mice received IT neopeptide vaccination regimen administered weekly for a total of four doses, starting when tumor size reached approximately 50 mm^3^ (the IT vaccination schedule shown is representative). **b**, Representative images of ulcers in tumors in the SQ saline + IT saline, SQ vaccine + IT adjuvants, and P/T vaccination groups (left). Ulcer diameters indicated by arrow. Quantification of ulcer diameter and ulcer frequency (right, n = 11-12 tumors per group). **c-d**, Average tumor growth curve (**c**) and Kaplan-Meier survival curve (**d**) of AE17 tumor-bearing mice treated as indicated. **e**, Fold change in tumor size relative to the tumor size at the time of the 1^st^ IT injection in the AE17 tumor model for individual mice shown. The data shown represent the time course post-1^st^ IT injection. Inverted triangles indicate IT injection time points. Data in **c-e** are from three independent experiments (n = 12-22 mice per group). **f**, Experimental design for P/T vaccination in the KPC-4662 tumor model. C57BL/6J mice were inoculated with KPC-4662 tumor cells (SQ, 0.3 x 10^6^ cells) on day 0. SQ neopeptide vaccines were administered on days 0, 7, 14, 28, and 42. A 5^th^ SQ vaccination on day 42 was performed only in the long-term experiment **(h-j)**. Tumor-bearing mice received IT neopeptide vaccination weekly for a total of four doses, starting when tumor size reached approximately 100 mm^3^ (the IT vaccination schedule shown is representative). **g**, Average tumor growth curve (left) and Kaplan-Meier survival curve (right) of KPC-4662 tumor-bearing mice treated with SQ saline + IT saline or P/T vaccination from two independent experiments (n = 15 or 17 mice per group). **h**-**i**, Average tumor growth curve (**h**) and Kaplan-Meier survival curve (**i**) of KPC-4662 tumor-bearing mice treated as indicated and monitored for long term. **j**, Fold change in tumor size relative to the tumor size at the time of the 1^st^ IT injection in the KPC-4662 tumor model for individual mice shown. The data shown represent the time course post-1^st^ IT injection. Inverted triangles indicate IT injection time points. Data in **h**-**j** are from a single experiment (n = 3-7 mice per group). Data represent the mean ± SEM (**b** and **g**) and were statistically analyzed using Kruskal-Wallis test with Dunn’s multiple comparisons test for the ulcer size, or Fisher’s exact test for the frequencies (**b**). Average tumor growth curves were statistically analyzed by two-way ANOVA with Bonferroni multiple comparisons test as of the end point of the SQ saline + IT saline group (**c**, **g**, and **h**). Survival curves were statistically analyzed by log-rank (Mantel-Cox) test (**d**, **g**, and **i**). \**p* < 0.05; \*\**p* < 0.01; \*\*\**p* < 0.001; \*\*\*\**p* < 0.0001.

### P/T neopeptide vaccination boosts tumor-infiltrating Th1 cells and non-exhausted effector CD8^+^ T cells

To gain insights into the cellular mechanism behind the effect of P/T neopeptide vaccination, we analyzed tumor-infiltrating T cells from mice that were primed by SQ neopeptide vaccination and then received a single IT neopeptide vaccination. Tumors were dissected 7 days post IT vaccination and analyzed by spectral flow cytometry (Fig. 2a). These experiments revealed that following P/T neopeptide vaccination, large numbers of T cells were recruited into tumors, the majority of which were CD4^+^ T cells (Fig. 2b, c). Consistent with their limited effects on tumor control in the AE17 tumor model (Fig. 1c-e), such T cell recruitment was not detected following SQ or IT vaccinations alone or SQ vaccination followed by IT adjuvant injection (Extended Data Fig. 2b). After P/T neopeptide vaccination, both the frequency and the absolute number of Tbet^+^ Th1 CD4^+^ T cells drastically increased in the tumors (Fig. 2d-g). This Th1 cell abundance increased over time following the IT vaccination, indicating enhanced recruitment and/or local proliferation of these cells within tumors (Extended Data Fig. 2c). By contrast, Treg cells, which suppress anti-tumor immune responses, were decreased upon P/T neopeptide vaccination compared to that of the saline group (Fig. 2d-g). Interestingly, P/T neopeptide vaccination did not affect the numbers of total CD8⁺ T cells within tumors (Fig. 2f, g) although their abundance trended to increase following IT vaccination (Extended Data Fig. 2c). In line with the flow cytometric data, immunofluorescence staining revealed abundant CD4⁺ T cell infiltration within the tumors 7 days after IT vaccination (Fig. 2h). These CD4⁺ T cells were distributed throughout the tumor parenchyma without clear regional accumulation at this time point.

**Fig. 2.**
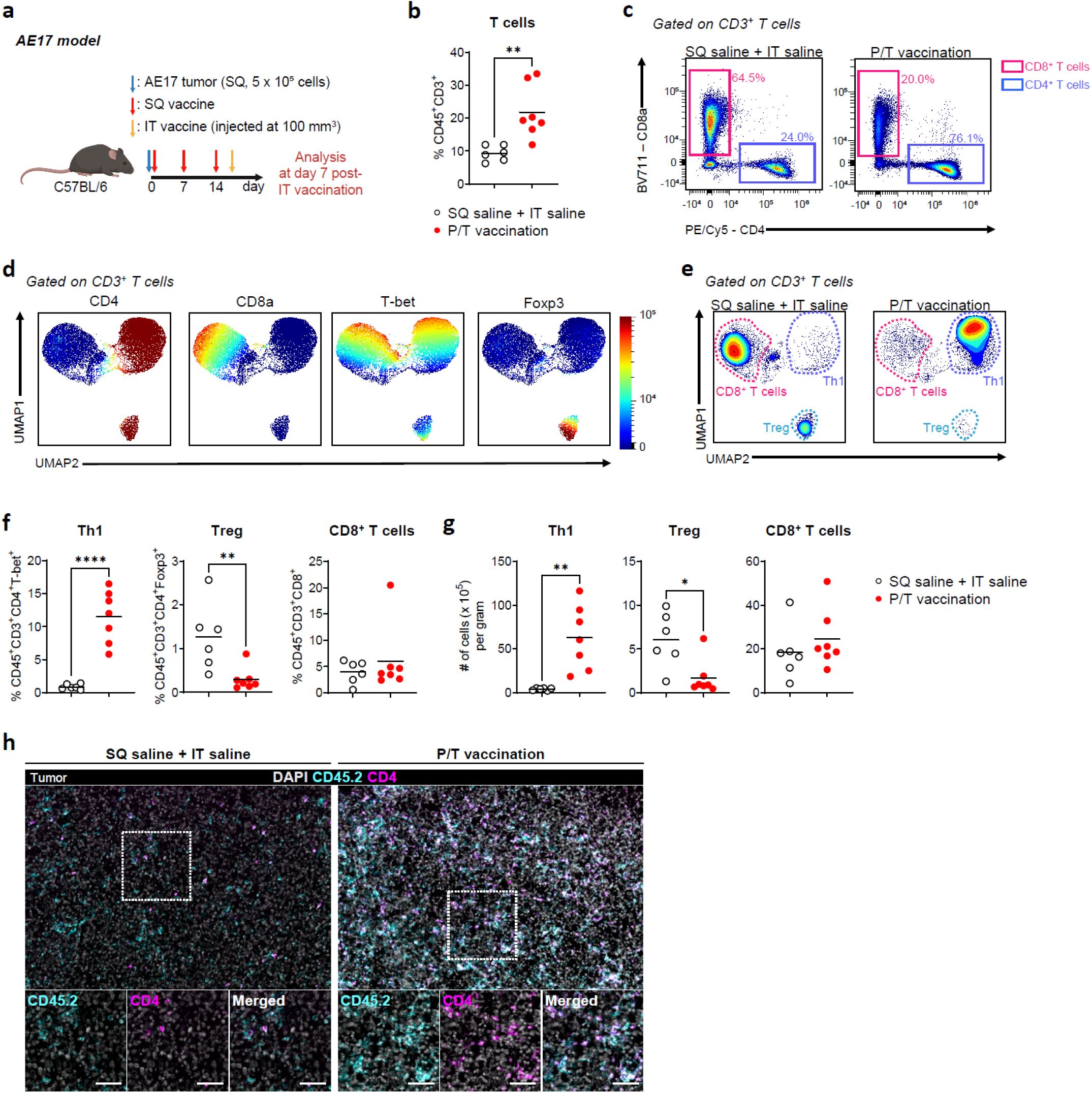
P/T neopeptide vaccination recruits massive numbers of CD4^+^ Th1 cells into tumors 7 days after a single IT vaccination. a, Experimental design to examine the TME following P/T vaccination in the AE17 tumor model. C57BL/6J mice were inoculated with AE17 tumor cells (SQ, 0.5 x 10^6^ cells) on day 0. SQ neopeptide vaccines were administered on days 0, 7 and 14. Meanwhile, tumor-bearing mice received an IT neopeptide vaccination once when tumor size reached approximately 100 mm^3^ (the IT vaccine schedule shown is representative). Tumor tissues were harvested 7 days after a single IT vaccination. **b,** Frequency of tumor-infiltrating T cells, defined as CD45^+^ CD3^+^ among live singlets, per group at day 7 after a single IT vaccination. **c,** Flow cytometric gating strategy for tumor-infiltrating T cells, showing CD4^+^ and CD8^+^ T populations (representative tumor samples shown). **d,** UMAPs of tumor-infiltrating T cells showing expression level of the indicated markers based on flow cytometric analysis (data pooled from 2 or 4 tumors per group). **e,** UMAPs of tumor-infiltrating T cells highlighting Th1 (CD45^+^ CD3^+^ CD4^+^ T-bet^+^), Treg (CD45^+^ CD3^+^ CD4^+^ Foxp3^+^), and CD8^+^ T cell (CD45^+^ CD3^+^ CD8^+^) subsets (representative tumor samples shown). **f-g,** Frequencies and absolute numbers per gram of tumor tissue of Th1, Treg, and CD8^+^ T cells. **h,** Immunofluorescence staining of tumor tissues for DAPI, CD45.2, CD4 (representative tumors shown). CD4^+^ cells were defined as CD45.2^+^CD4^+^. Scale bar, 50 µm. Data in b-h are from two independent experiments (n = 6 or 7 mice per group). Horizontal lines in b, f, and g represent the mean and were statistically analyzed by two-tailed unpaired Student’s t-test or two-tailed unpaired Mann-Whitney U-test. \**p* < 0.05; \*\**p* < 0.01; \*\*\**p* < 0.001; \*\*\*\**p* < 0.0001.

To gain further insight into the changes P/T neopeptide vaccination induced in tumor-infiltrating T cells and the TME, we performed single-cell RNA sequencing (scRNA-seq) of tumor tissues 7 days after a single IT vaccination of SQ primed animals (Fig. 2a). Single-cell suspensions of tumors were labeled with TotalSeq antibody-derived tags (ADT), and live cells were subsequently sorted by flow cytometry and subjected to scRNAseq. Clustering of *Ptprc*⁺ (CD45⁺) immune cells based on ADT protein expression revealed five metaclusters (Extended Data Fig. 3a). Subsequently, metaclusters 1 and 2, which were positive for CD4 or CD8α at both the RNA and protein levels (Extended Data Fig. 3b, c), were analyzed as T cell cluster. Based on canonical marker expression and the top 10 differentially expressed genes (DEGs), the T cell cluster was further categorized into 11 T-cell subclusters, all *Cd3e*-positive: five CD4^+^ Th1 cell (CD4_Th1) subclusters, one CD4^+^ Treg cell (CD4_Treg) subcluster, one effector CD8^+^ T cell (CD8_Eff) subcluster, and three exhausted CD8^+^ T cell subclusters including terminally exhausted (CD8_TEX-term), proliferating exhausted (CD8_TEX-prolif), and progenitor-like exhausted (CD8_TPEX) subclusters, as well as one mixed CD4/CD8 subcluster containing both CD4⁺ and CD8⁺ T cells (Fig. 3a, b and Extended Data Fig. 4). All of the 11 T-cell subclusters were present in both the control (SQ saline + IT saline) and P/T neopeptide vaccination groups, however, their distribution was greatly altered by P/T neopeptide vaccination (Fig. 3c). Consistent with the finding in flow cytometric analysis (Fig. 2e-g), tumor-infiltrating T cells were predominantly Th1 cells in tumors from mice that received P/T neopeptide vaccination (Fig. 3c). In particular, activated Th1 cell subsets expressing *Cd40lg* (CD4_Th1_CD40L+), *Tnfrsf4* (CD4_Th1_OX40+), or interferon-stimulated genes (ISGs) such as *Isg15* and *Isg20* (CD4_Th1_ISG+) were significantly increased (Fig. 3d, e). While exhausted CD8⁺ T cell subsets were not altered by the P/T neopeptide vaccination, effector CD8⁺ T cells tended to increase (Fig. 3d, e). These finding suggests that P/T neopeptide vaccination can thwart tumor- mediated T cell exclusion or suppression and promotes T cells with effector potential within the TME.

**Fig. 3.**
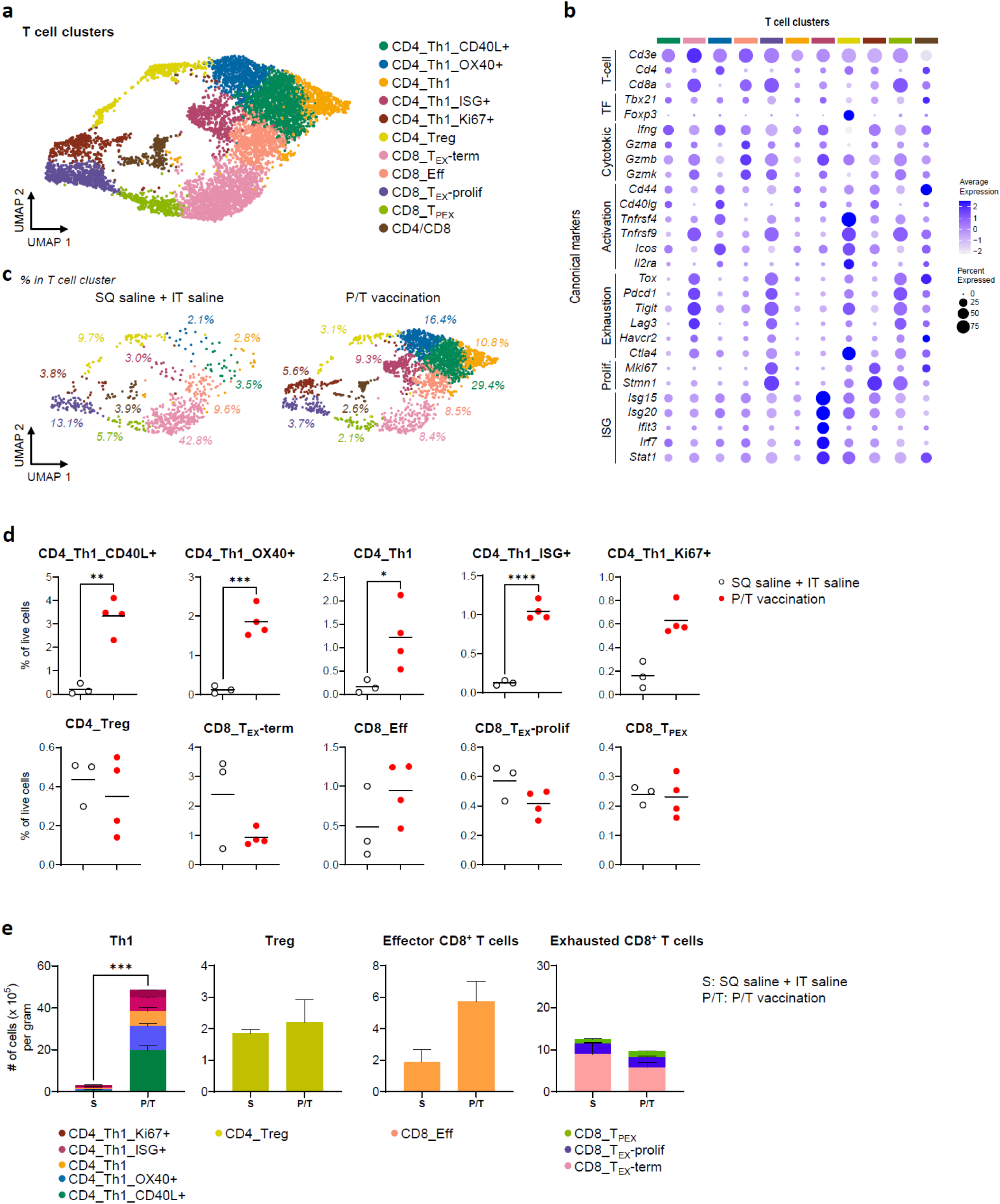
Tumor-infiltrating Th1 following P/T neopeptide vaccination exhibit activated phenotypes. Animals were treated as in Fig. 2a. **a**, UMAP of scRNA-seq of the T cell cluster with cell types in tumors at day 7 after a single IT vaccination. scRNA-seq was performed on FACS-sorted live cells from digested tumors. **b**, Dot plot of canonical markers identifying CD4^+^ Th1, CD4^+^ Treg, and CD8^+^ T cell subsets. **c**, UMAPs of the T cell cluster in SQ saline + IT saline (left, n = 3 mice) and P/T vaccination (right, n = 4 mice). **d**, Frequencies of the T cell subclusters among total live cells of tumors. **e**, Absolute numbers of the T cell subclusters per gram of tumor tissue. Data in **a**-**e** represent two separate sequencing runs conducted across two independent experiments (n = 3 or 4 per group). Horizontal lines in **d** indicate the mean. Data were statistically analyzed by ordinary one-way ANOVA with Tukey’s multiple comparisons test or Kruskal-Wallis test with Dunn’s multiple comparisons test. Bars in **e** represent the mean ± SEM, and stacked subsets were statistically analyzed by two-tailed unpaired Student’s t-test . \**p* < 0.05; \*\**p* < 0.01; \*\*\**p* < 0.001; \*\*\*\**p* < 0.0001.

### P/T neopeptide vaccination induces large clonal expansions of neoantigen-specific Th1 cells and effector CD8^+^ T cells in tumors

Given the striking changes in T cell populations observed within the TME, we next asked whether P/T neopeptide vaccination also reshaped the TCR repertoire of tumor-infiltrating T cells. We therefore analyzed single-cell T cell receptor sequencing (scTCR-seq) data generated in parallel to scRNA-seq and examined complementarity-determining region 3 (CDR3) sequences, which determine the antigen-specificity of T cells and enable tracking of clonal expansion. Overall, we found increased TCR diversity with many more CDR3 motifs in tumors after P/T neopeptide vaccination (Fig. 4a), most of which were not present in SQ+IT saline-injected controls. In contrast, one motif (motif 8 CASIG.QNTLYF), was broadly shared across animals of both groups, indicating that this clonotype likely represents endogenous tumor-reactive T cells (Fig. 4a, b). Of the top 30 CDR3 motifs, most were dominantly detected in Th1 cells (Fig. 4a). Notably, more Th1-associated clones were observed and shared in the P/T-vaccinated mice suggesting that Th1 cells undergo dominant clonal expansion following P/T neopeptide vaccination (Fig. 4a, b). Consistent with this, the Gini index of Th1 cells, an indicator of clonality, was significantly higher in the P/T neopeptide vaccination group compared with the control group (Fig. 4c), indicating increased clonal dominance and reduced repertoire evenness following P/T neopeptide vaccination. Furthermore, effector CD8^+^ T cells in the P/T neopeptide vaccination group also showed greater clonal expansion, whereas exhausted CD8⁺ T cells remained unchanged and had as dominant clone the TCR with motif 8 mentioned above (Fig. 4a-c). This clone was shared between exhausted CD8^+^ T cells of control and P/T vaccinated animals (Fig. 4a, b). Consistently, highly expanded T cell clones with large clone sizes were predominantly detected in Th1 and effector CD8⁺ T cell subsets of the P/T neopeptide vaccination group (Fig. 4d, e). Taken together, these findings indicate that P/T neopeptide vaccination drives clonal expansion of Th1 and effector CD8⁺ T cells without altering the pre-existing exhausted CD8⁺ T-cell compartment.

**Fig. 4.**
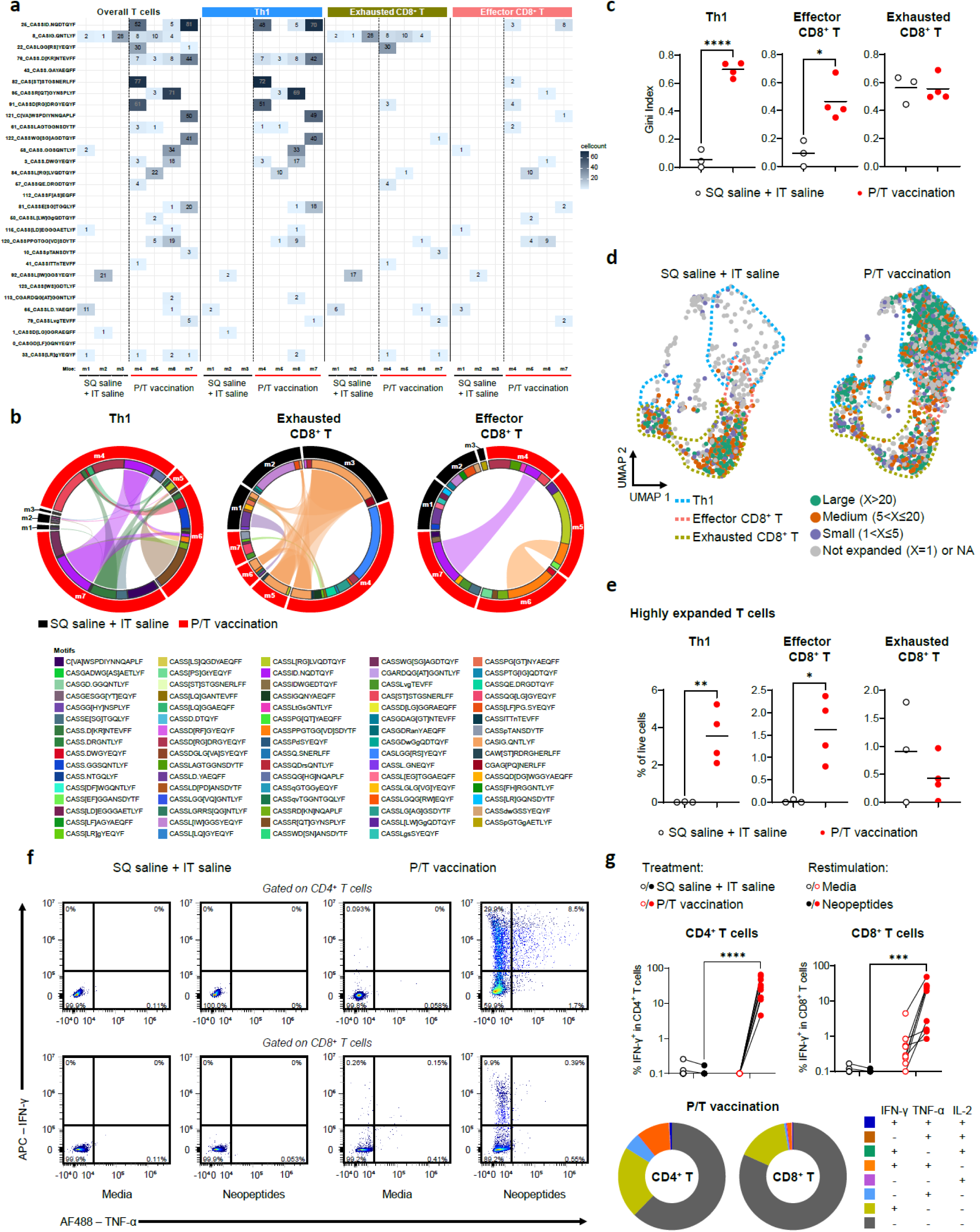
P/T neopeptide vaccination recruits into tumors vaccine-induced Th1 and effector CD8+ T cells harboring newly emerging TCR clones. Animals were treated as in Fig. 2a. Cells from the scRNAseq T cell clusters were analyzed based on their TCRβ CDR3 amino acid sequence (tumors at day 7 after a single IT vaccination). The top 30 CDR3 motifs were selected according to their frequency among all clustered T cells across all samples and groups. **a,** Heatmap showing cell counts for the top 30 CDR3 motifs across tumor-infiltrating overall T cells or within individual pooled T cell subclusters in the SQ saline + IT saline and P/T vaccination groups. Pooled Th1 cell subcluster includes CD4_Th1, CD4_Th1_CD40L+, CD4_Th1_OX40+, CD4_Th1_ISG+, and CD4_Th1_Ki67+ subsets; pooled exhausted CD8^+^ T cell subcluster includes CD8_TEX-term, CD8_TEX-prolif, and CD8_TPEX subsets; effector CD8+ T cells correspond to CD8_Eff subset. Each column represents an individual mouse (n = 3 or 4 mice per group). **b,** Circos plots illustrating shared T cell clones based on all identified CDR3 motifs across pooled Th1, pooled exhausted CD8+ T cell, and effector CD8+ T cell subclusters in the SQ saline + IT saline (black arcs, mice m1, m2 and m3) and P/T vaccination (red arcs, mice m4, m5, m6 and m7) groups. Individual mice are shown (m1 to m3, SQ saline + IT saline; m4 to m7, P/T vaccination). Each CDR3 motif is color-coded and connecting lines indicate shared motifs between mice. **c,** Gini index of individual mice for pooled Th1, pooled exhausted CD8^+^ T cell and effector CD8^+^ T cell subsets in tumors shown (n = 3 or 4 mice per group). **d,** UMAPs showing expanded clones, colored based on clone size, in SQ saline + IT saline (left, n = 3 mice) and P/T vaccination (right, n = 4 mice) in tumors. Each dot depicts an individual T cell. Grey dots indicate unexpanded single clones (X = 1), purple dots indicate small size clones (shared by 1 < X ≤ 5 cells), brown dots indicate medium size clones (shared by 5 < X ≤ 20 cells), and green dots indicate large size clones (shared by X>20 cells). **e,** Frequencies of highly expanded T cells with large size clones (X>20 cells in tumor) shown in Fig. 4d in pooled Th1, pooled exhausted CD8^+^ T cell, and effector CD8^+^ T cell subsets in tumors (n = 3 or 4 mice per group). **f,** Neopeptide-specific T cells in tumors 7 days after IT vaccination. Representative flow cytometric plots showing neopeptide-specific T cells producing IFN-γ and/or TNF-α shown. Neopeptide-specific cytokine production was assessed upon restimulation with the corresponding neopeptides; media condition served as a negative control. **g,** Top, frequencies of neopeptide-specific IFN-γ-producing CD4^+^ or CD8^+^ T cells in the SQ saline + IT saline and P/T vaccination group (n = 9 mice per group from three independent experiments). Bottom, distribution of cytokine-producing (IFN-γ, TNF-α, and/or IL-2^+^) cells within tumor-infiltrating CD4^+^ and CD8^+^ T cells in the P/T vaccination group (n = 9, average). Horizontal lines in c and e represent the mean and data were statistically analyzed by two-tailed unpaired Student’s t-test or two-tailed unpaired Mann-Whitney U-test. Data in **g** were statistically analyzed by two-way ANOVA with Bonferroni multiple comparisons test. \**p* < 0.05; \*\**p* < 0.01; \*\*\**p* < 0.001; \*\*\*\**p* < 0.0001.

We next examined whether tumor-infiltrating T cells following P/T neopeptide vaccination were specific for neopeptides included in our vaccine formulation. To this end, digested tumors were restimulated with vaccine neopeptides, and cytokine production by T cells was assessed by flow cytometry directly *ex vivo*. The murine H-2^b^ DC cell line DC2.4 was added during restimulation to provide sufficient antigen cross-presentation of 20mer neopeptide. These experiments showed that P/T neopeptide vaccination vastly increased the frequency of neopeptide-specific T cells within the TME compared to the control group receiving SQ and IT saline (Fig. 4f, g). On average, 37.6% of CD4⁺ T cells and 18.1% of CD8⁺ T cells were neopeptide-specific, and the majority of these cells produced IFN-γ alone followed by cells co-producing IFN-γ and TNF-α. The frequencies of neopeptide-specific T cells within the TME following P/T neopeptide vaccination were higher than in the spleens of SQ-vaccinated mice (Extended Data Fig. 1a), indicating enhanced recruitment and/or expansion of vaccine-induced T cells within the TME following P/T neopeptide vaccination. In contrast, the frequency of neopeptide-specific T cells, particularly CD8^+^ T cells, were markedly less abundant in the tumor-draining lymph nodes (tdLNs) compared to the tumors at day 7 (Extended Data Fig. 5a). These data indicate that neopeptide-specific T cells recruited following P/T vaccination tended to undergo preferential expansion within tumors, with limited expansion in tdLNs. Consistently, in tumors Ki67⁺ Th1 cells and non-exhausted (PD-1⁻) CD8⁺ T cells increased over time and were more frequent than in control mice (Extended Data Fig. 5b), suggesting active proliferation of tumor-infiltrating T cells. Collectively, these findings demonstrate that P/T neopeptide vaccination predominantly recruits and expands clonal populations of neopeptide-specific Th1 and effector CD8⁺ T cells locally within tumors.

### Reduced M2-like macrophages and increased type-I IFN signature in the TME after P/T neopeptide vaccination

As priming of T cells requires antigen-presenting myeloid cells, such as DCs, for their activation in cancer immunity^24,25^, but also for their expansion within effector sites^19^, we additionally analyzed major myeloid subsets including monocytes, DCs, and tumor-associated macrophages (TAM) in the TME by flow cytometry 7 days after a single IT vaccination following systemic SQ vaccination (Fig. 2a and Extended Data Fig. 6a). None of these populations, except for the TAM population, were significantly altered following P/T neopeptide vaccination (Extended Data Fig. 6b). To more deeply profile possible changes in this compartment, we examined the transcriptional profiles of myeloid cells derived from our scRNA-seq data (metaclusters 0, 3, and 4; Myeloid cluster), which were positive for *Itgam* (CD11b), *Itgax* (CD11c), and/or *Adgre1* (F4/80) (Extended Data Fig. 3a-c). Based on their canonical marker expression and the top 10 DEGs, this Myeloid cluster was further categorized into 12 Myeloid subclusters: one monocyte-derived DC subset (Mo-DC), five TAM subsets (TAM_1, TAM_2, TAM_3, TAM_4, TAM_5), of which TAM_1 to TAM_4 exhibit a clear M2 phenotype ^26–28^, two monocyte subsets (Mono_1, Mono_2), three DC subsets (cDC1, cDC2, mature DC enriched in immunoregulatory molecules (mregDC)), and one myeloid-derived suppressor cell subset (MDSC) (Fig. 5a, b and Extended Data Fig. 7a). Consistent with the finding of our flow cytometric analysis (Extended Data Fig. 6b), P/T neopeptide vaccination reduced the composition of TAM subsets and particularly the absolute number of TAM_1 and TAM_2 populations, which predominantly exhibited an M2-like phenotype characterized by expression of *Arg1* and *Mrc1* (Fig. 5c, d and Extended Data Fig. 7b). Besides M2-like TAM reduction, monocyte populations, especially Mono_2, were significantly increased following P/T vaccination (Fig. 5c, d and Extended Data Fig. 7b).

**Fig. 5.**
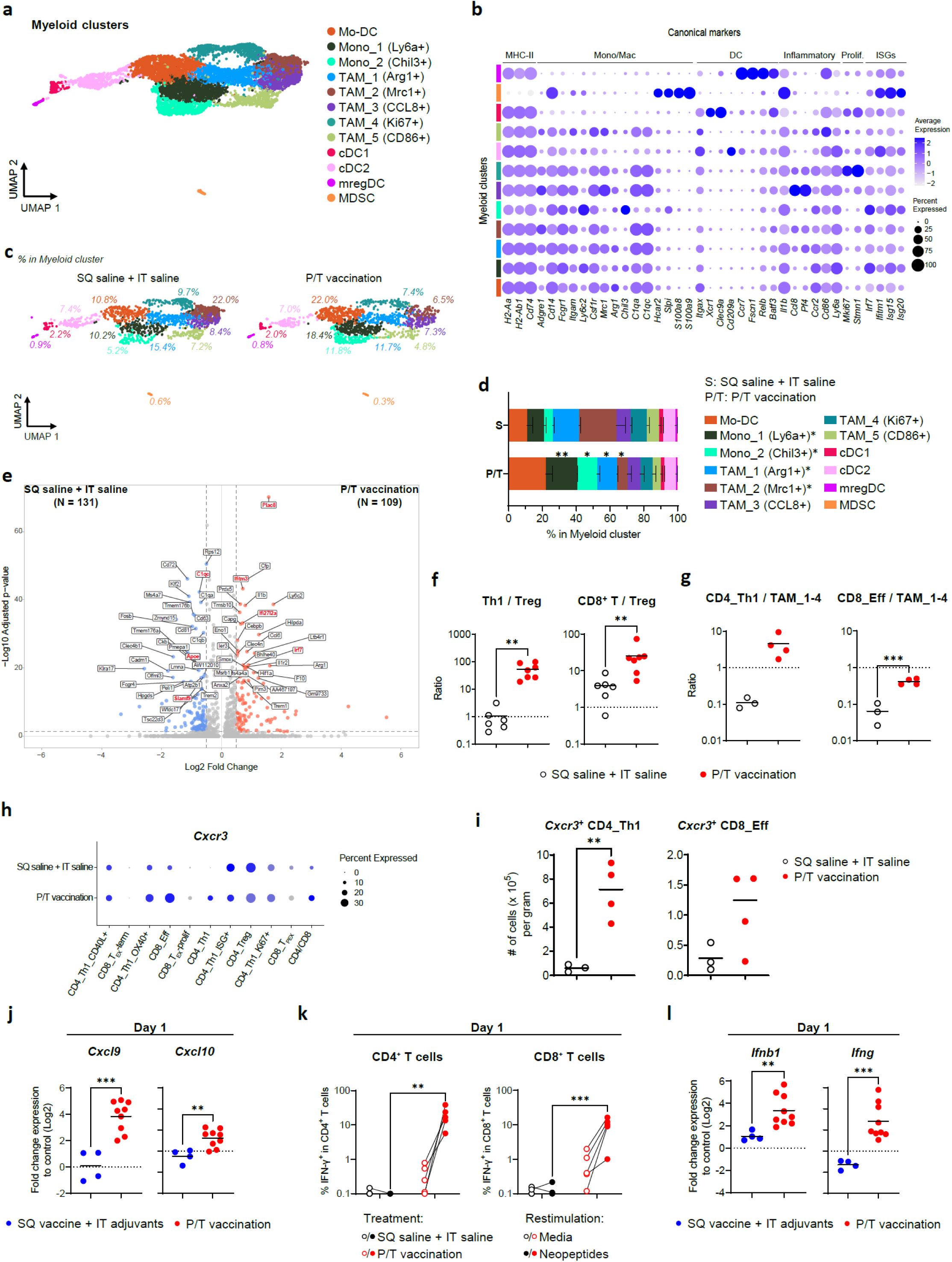
P/T neopeptide vaccination remodels the TME toward an anti-tumor phenotype. **a**, UMAP of scRNA-seq of the myeloid cluster with cell types in tumors at day 7 after a single IT vaccination. See Extended Data Fig. 3a fig for identification of the myeloid cluster. **b**, Dot plot of canonical markers identifying monocyte, macrophage, and DC subsets. **c**, UMAPs of the myeloid cluster in the SQ saline + IT saline (left, n = 3 mice) and P/T vaccination (right, n = 4 mice) groups. **d**, Composition of myeloid cell subsets within the myeloid cluster in the SQ saline + IT saline and P/T vaccination groups. Colors in bars correspond to populations in a. **e**, Volcano plots showing differentially expressed genes in the myeloid cluster between the SQ saline + IT saline and P/T vaccination groups. Statistically significant genes are colored in blue (high in SQ saline + IT saline) or red (high in P/T vaccination). **f**, Ratio of effector T cells (Th1 (left), CD8**^+^**T cells (right)), and Treg in tumors at 7 days after a single IT vaccination, based on flow cytometric analysis shown in Fig. 2f. **g**, Ratio of effector T cells (pooled Th1 subsets (left), effector CD8**^+^** T cell subsets (CD8_Eff, right)), and M2-like macrophages (pooled TAM_1 to _4) in tumors at 7 days after a single IT vaccination, based on scRNA-seq analysis shown in Fig. 3a and 4a. **h**, Dot plot showing *Cxcr3* expression levels in individual T cell subsets within tumors from the SQ saline + IT saline and P/T vaccination groups (n =3 or 4 per group, average), analyzed by scRNA-seq shown in Fig. 3a. **i**, Absolute numbers of Cxcr3-expressing pooled Th1 (left) and effector CD8+ T cell (right) subsets per gram of tumor tissue in the SQ saline + IT saline and P/T vaccination groups (n =3 or 4 per group), analyzed by scRNA-seq shown in Fig. 3a. **j**, Fold change in *Cxcl9* (left) and *Cxcl10* (right) expression, analyzed by qPCR, in tumors harvested one day after a single IT vaccination in the SQ vaccine + IT adjuvants (blue) and P/T vaccination (red) primed groups (n= 4 or 9 per group from three independent experiments). Fold change was normalized to the SQ saline + IT saline group and converted to Log2 scale. **k**, Frequencies of neopeptide-specific CD4**^+^** or CD8+T cells in the tumors harvested one day after a single IT vaccination. SQ saline + IT saline and P/T vaccination groups shown (n = 4 or 5 mice per group, one experiment). Neopeptide-specific cytokine production was assessed upon restimulation with the corresponding neopeptides; media condition served as a negative control. **l**, Fold change in *Ifnbl* (left) and *ifng* (right) expression, analyzed by qPCR, in tumors harvested one day after a single IT vaccination in the SQ vaccine + IT adjuvants and P/T vaccination groups (n= 4 or 9 per group from three independent experiments). Fold change was normalized to the SQ saline + IT saline group and converted to Log2 scale. Data in **a**-**e** and **g**-**i** are from two separate sequencing runs conducted across two independent experiments (n = 3 or 4 tumors per group). Data in **d** represent the mean **±** SEM were statistically analyzed by two-tailed unpaired Student’s t-test. Horizontal lines in **f**, **g**, **i**, **j**, and **l** represent the mean. Data were statistically analyzed by two-tailed unpaired Student’s t-test or two-tailed unpaired Mann-Whitney U-test. Data in **k** were statistically analyzed by two-way ANOVA with Bonferroni multiple comparisons test. \**p* < 0.05; \*\**p* < 0.01; \*\*\**p* < 0.001; \*\*\*\**p* < 0.0001.

To further dissect signaling within the myeloid compartment after P/T neopeptide vaccination, we analyzed DEGs in the entire myeloid cluster. Among the significantly upregulated genes by P/T neopeptide vaccination, several were associated with type-I IFN signaling, including *Irf7*, *Plac8*, *Ifitm3*, and *Ifi27l2a* (Fig. 5e). Consistent with these findings, gene ontology (GO) analysis of the pseudo-bulk myeloid cluster revealed that P/T neopeptide vaccination strongly upregulated pathways associated with IFN responses, including positive regulation of cytokine production and negative regulation of viral genome replication pathways (Extended Data Fig. 7c). Conversely, P/T neopeptide vaccination downregulated genes linked to immunosuppression, such as *C1qc*, *Apoe*, and *Slamf9* (Fig. 5e). Together, the above results indicated that P/T neopeptide vaccination reduced subsets of M2-like macrophages within the TME while the entire myeloid compartment adopts a type I IFN-associated activated phenotype. Such TME M2 population changes are consistent with less suppression and improved anti-tumor activity by infiltrating T cells.

### P/T neopeptide vaccination drastically remodels the TME toward an anti-tumor phenotype

The observed reduction in immunosuppressive cells, such as Treg (Fig. 2f, g) and certain subsets of M2 macrophages (Fig. 5c, d and Extended Data Fig. 7b), by P/T vaccination prompted us to further evaluate the microenvironment encountered by T cells within the TME. To this end, we quantified the ratio of effector T cells to either Treg cells or M2 macrophages using either the flow cytometric or scRNA-seq data, as these balances are critical determinants of anti-tumor immunity^29,30^. P/T neopeptide vaccination markedly increased the Th1/Treg, CD8⁺ T cell/Treg, Th1/M2-like macrophage and effector CD8⁺ T cell/M2-like macrophage ratios (Fig. 5f, g), with the latter calculated using pooled M2-like clusters TAM_1 to TAM_4. Together, these changes indicate that P/T neopeptide vaccination not only enhances effector T-cell infiltration and proliferation, but also reshapes the myeloid and regulatory landscape to favor effective T cell immunity within tumors.

### P/T vaccination enables rapid T cell recruitment into tumors

In our tumor scRNA-seq data, Th1 subsets and effector CD8⁺ T cell subset, but not exhausted CD8⁺ T cell subsets, expressed *Cxcr3*, a key chemokine receptor required for T cell infiltration into tumors^31,32^ (Fig. 5h, i). We therefore examined by qPCR whether P/T neopeptide vaccination induced expression of the CXCR3 chemokine ligands *Cxcl9* (MIG) or *Cxcl10* (IP-10) within tumors to seek a possible mechanism of T cell recruitment into tumors. Indeed, P/T neopeptide vaccination significantly upregulated *Cxcl9* and *Cxcl10* expression at day 1 post IT vaccination (Fig. 5j) although these chemokines were no longer elevated on day 7, when T-cell infiltration peaked (Extended Data Fig. 7d and 2c). This suggested that a subset of systemically primed, vaccine-induced T cells may begin entering tumors as early as day 1 after IT vaccination. To test this, we analyzed tumor-infiltrating neopeptide-specific T cells following *ex vivo* antigen restimulation one day after a single IT vaccination (mice were primed by SQ neopeptide vaccination). As anticipated, both neopeptide-specific CD4⁺ and CD8⁺ T cells were already prominently increased in P/T-vaccinated tumors one day after IT vaccination (Fig. 5k). Consistent with the early recruitment of neopeptide-specific T cells capable of producing IFN-γ upon antigen engagement, and given that IFN-γ and, to a lesser extent, type-I IFN are known inducers of CXCL9/10^33^, we examined by qPCR whether IFNs were produced in tumors at this initial time point. Both *Ifnb1* and *Ifng*, which indicate a pro-inflammatory and anti-tumor TME phenotype^34,35^, were markedly upregulated by P/T neopeptide vaccination one day after a single IT vaccination (Fig. 5l). This upregulation was not exclusively due to the presence of adjuvants, as SQ vaccination followed by IT injection only with adjuvants exhibited just a weak induction of IFN-β and no IFN-γ (Fig. 5l). Therefore, IT antigen plus adjuvants were both required for the early upregulation of IFNs in the TME. These findings indicate that P/T neopeptide vaccination initiates a rapid positive feedback amplification loop of IFNs, chemokines and T cell recruitment in tumors that supports additional waves of neopeptide-specific T-cell recruitment, ultimately enabling robust T cell infiltration and expansion.

## DISCUSSION

Here, we report a novel P/T neopeptide vaccination strategy that first induces strong systemic neoantigen-specific T cell immunity in the periphery and then – via IT vaccination – recruits and expands large numbers of neoantigen-specific Th1 and non-exhausted effector CD8^+^ T cells within tumors. This unique and novel vaccination strategy exhibits unprecedented control and cures of established immunotherapy-resistant mouse tumors, such as mesothelioma and pancreatic adenocarcinoma. This protection was accompanied by a striking immunological rewiring of the TME that was characterized by strong expansion of activated Th1 cells and non-exhausted effector CD8^+^ T cells within tumors while suppressive Treg and M2 macrophages were reduced. Efficacy of P/T vaccination depended on neopeptide and adjuvant combination being used for both SQ priming and IT vaccination as SQ priming followed by IT injection with adjuvants alone or neopeptides alone failed to protect. Thus, it is critical to concurrently deliver locally both antigen and adjuvants to harvest the benefit of potent systemic T cell priming.

Solid tumors rich in T cells, which are often referred to as “hot” tumors, are typically more responsive to cancer immunotherapies, whereas immunosuppressive “cold” tumors with sparse T cell infiltration frequently fail to respond^3,4^. Therefore, converting a “cold” TME into a T cell-inflamed and pro-inflammatory state is a critical step toward achieving effective cancer immunotherapy^1,8,9^. Here, we demonstrate that P/T neopeptide vaccination achieves two complementary goals: (1) priming strong systemic anti-tumor T cell responses through SQ vaccination, and (2) remodeling the TME to facilitate effective tumor cell targeting through IT vaccination. Specifically, P/T neopeptide vaccination potently increases the effector T cell compartment and reprograms the myeloid compartment toward an IFN-associated, activated phenotype. The induction of type-I IFNs and ISGs such as *Isg15* and *Isg20* in the tumors has previously been shown to be associated with improved outcomes in cancer immunotherapy^35-37^. Thus, the establishment of an IFN-associated pro-inflammatory TME following P/T neopeptide vaccination is likely critical for its therapeutic efficacy. Strikingly, neopeptide-specific CD4^+^ and CD8^+^ T cells and *Ifnb1* and *Ifng* expression in the tumors were detectable as early as one day after a single IT vaccination in P/T neopeptide-vaccinated mice, indicating that TME modulation by P/T neopeptide vaccination occurs rapidly and requires tumor-specific antigens, as *Ifng* was absent and *Ifnb1* only weakly induced by IT adjuvant administration in SQ primed animals. This strongly suggests that P/T neopeptide vaccination may be more effective than *in situ* vaccination strategies, that only utilize intratumoral administration of adjuvants like TLR3 or STING agonists, to transform the TME into a pro-inflammatory milieu ^10,21,22,38^. Based on our findings, we hypothesize that IT vaccination establishes a positive feedback loop in the tumor of T cell activation and recruitment. Based on this model, the initial activation of rare neoantigen-specific T cells in tumors would induce IFN-γ and amplify type I IFN production, both of which in turn induce the expression of CXCL9/CXCL10. This leads to further CXCR3^+^ neoantigen-specific T cell recruitment from circulation and more local cytokine production. This loop of intratumor T cell activation, T cell recruitment, cytokine and chemokine production requires cognate neoantigens included in the systemic and IT vaccination.

P/T neopeptide vaccination induced necrotic lesions and tumor shrinkage indicating tumor cell killing, however, the precise mechanism of how tumor cells are eliminated remains to be fully elucidated. The majority of tumor-infiltrating T cells induced by P/T neopeptide vaccination was specific for neopeptides contained in the vaccine formulation. Given the massive influx of Th1 cells after P/T vaccination, we hypothesize that these neopeptide-specific tumor-infiltrating Th1 cells are key in mediating the anti-tumor response. CD4^+^ effector Th1 cells are known to play a critical helper role in supporting effector CD8^+^ T cells in cancer immunity^6,8,24,39,40^. Accordingly, the Th1 cells recruited with our P/T neopeptide vaccination likely help to establish a type-1 immunity-prone microenvironment that is favorable for the recruitment, activation and persistence of cytotoxic CD8^+^ T cells. In line with this, neopeptide-specific CD4^+^ T cells have been shown to counteract the immunosuppressive TME, thereby promoting the induction of cytotoxic CD8^+^ T cells^41^, and this could underlie the marked expansion of non-exhausted effector CD8^+^ T cells we observed with P/T neopeptide vaccination. Such effector CD8^+^ T cells can directly mediate tumor killing. We cannot, however, exclude the possibility that neopeptide-specific Th1 cells in our system may directly kill tumor cells in a MHC-II-dependent manner, as suggested in recent studies^42–45^. Our scRNA-seq analysis revealed that the ISG^+^ Th1 population expresses *Gzmb* and *Gzmk*, but also *Tnfsf6* (encoding FasL) suggesting that this specific Th1 population may function as cytotoxic CD4^+^ T cells. Since baseline MHC-II is low to undetectable on our tumor models^46,47^, this would argue that MHC-II-expressing antigen presenting cells may trigger Th1 cells to rather deliver indirect killing of tumor cells in addition to helping CD8^+^ T cell-mediated direct tumor cell killing.

P/T neopeptide vaccination employs intra-tumor neopeptide vaccine delivery, which could be perceived as a limitation to its implementation in patient treatment. However, intratumor and intranodal delivery of immunotherapies or adjuvants are being tested already in many clinical trials^48^, indicating that intra-tumor delivery of a neopeptide vaccine would not represent an insurmountable obstacle. This together with its efficacy against immunotherapy-resistant tumors highlights P/T neopeptide vaccination as a viable option for patients with limited therapeutic options.

Overall, P/T neopeptide vaccination consistently produced robust anti-tumor efficacy across two hard-to-treat tumor models tested. We believe P/T vaccination need not be limited to neoantigens but could be employed with off-the-shelf targets such as tumor-associated antigens (TAA) and driver mutations, something that could enable wider application. Beyond its direct therapeutic potential, P/T vaccination could also be employed to facilitate the intratumoral recruitment of T cells from adoptive cell transfer (ACT) such as chimeric antigen receptor (CAR) T cells. Our approach can also serve as a tool to identify critical parameters that orchestrate TME remodeling in “cold” tumors and overcome T cell exclusion. Understanding such TME remodeling would be a major therapeutic breakthrough for cancers with limited treatment options. In conclusion, we demonstrate that our P/T neopeptide vaccination approach results in unprecedented protection and offers a promising therapeutic vaccination strategy for overcoming the resistance of immunosuppressive solid tumors to immunotherapy.

## METHODS

### Mice

Eight-week-old female C57BL/6J mice were purchased from Charles Rivers. Animal studies were carried out in accordance with the recommendations of the local authorities (Instantie voor Dierenwelzijn (IvD) that approved all protocols (license AVD1010020209604). Experimental animals were housed in the Erasmus Medical Center animal facility (Erasmus Dierenexperimenteel Center, EDC) in groups of five mice, food and water was administered *ad libitum*.

### Cell lines

AE17 mesothelioma cells (kindly gifted from Dr. D. Nelson, Curtin University, Perth, Australia) were cultured in RPMI media (Gibco) supplemented with 10% FBS (Gibco), 1% L-glutamine (Gibco), 50 µM β-mercaptoethanol (Sigma-Aldrich), 48 µg/ml gentamicin (Sigma-Aldrich) and 60 µg/ml benzylpenicillin (Qiagen). KPC-4662 pancreatic ductal adenocarcinoma cells (kindly gifted from Dr. R. H. Vonderheide, University of Pennsylvania, PA, USA) were cultured in DMEM media (Gibco) supplemented with 10% FBS, 1% L-glutamine, 100 units/ml Penicillin/Streptomycin (Gibco). DC2.4 cell line (initially contributed by K. Rock, University of Massachusetts Medical School, Worcester, Massachusetts, USA.) was cultured in RPMI media supplemented with 10% FBS, 1% HEPES (Gibco), and 100 units/ml Penicillin/Streptomycin. All cell lines were cultured at 37°C and 5% CO2.

### Neopeptides and adjuvants

Neoantigens for the AE17 and KPC-4662 tumors (listed in Supplementary Table 1) were identified using an in-house proprietary bioinformatic prediction pipeline as described previously^18^. Briefly, tumor-specific mutations in each tumor cell line were detected by whole-exome sequencing (WES), using C57BL/6 spleens as a reference, and RNA-seq was performed to assess the expression levels of mutated genes. Personalized Variant Antigens by Cancer Sequencing (pVAC-Seq) was utilized to integrate data of tumor mutations and expressions without applying the epitope predictor^49^. Identified neoantigens were synthesized as 20-mer neopeptides at Genscript and administered at 7 µg per peptide per vaccination for both SQ and IT vaccinations. For adjuvants, the TLR9 agonist K-type CpG oligodeoxynucleotide (K3 CpG) and the STING agonist c-di-AMP adjuvants were administered in combination (K3/c-di-AMP) at 10 µg each per vaccination for both SQ and IT vaccinations. K3 CpG was synthesized by GeneDesign (Japan) and c-di-AMP was kindly provided by Yamasa (Japan).

### In vivo tumor models

A frozen vial of each tumor cell line was thawed and cultured with two or three passages in appropriate culture media prior to tumor inoculation. For tumor inoculation, cells were harvested using 0.05% trypsin-EDTA (Gibco), washed with HBSS (Corning Life Sciences), resuspended in saline (Thermo Fisher Scientific), and mixed 1:1 (v/v) with Matrigel (Corning Life Sciences). Mice were anesthetized with 2-4% isoflurane and subcutaneously injected in the left flank with 100 µl of the tumor cell suspension containing 0.5 × 10^6^ tumor cells for the AE17 model or 0.3 × 10^6^ tumor cells for the KPC-4662 model. Mice were monitored for tumor growth and survival. Tumor size was measured three times a week using a digital vernier caliper, and tumor volume was calculated according to the following formula: volume = length (longest diameter) × width (shortest diameter) × height. Mice were euthanized when tumor volume reached the humane endpoint of 1,500 mm³.

### Vaccinations

To evaluate the therapeutic efficacy of the P/T neopeptide vaccination in the AE17 and KPC-4662 tumor models, tumor-bearing mice received 100 µl of SQ neopeptide vaccine containing corresponding 20-mer neopeptides and K3/c-di-AMP in the right flank on days 0, 7, 14, 28, and 42 post-tumor inoculation. For IT vaccinations, when tumor volume reached 50 or 100 mm^3^ size, tumor-bearing mice received 25-30µl (AE17) or 35µl (KPC-4662) of IT neopeptide vaccine containing corresponding 20-mer neopeptides and K3/c-di-AMP weekly, up to four times. For P/T vaccinations using pathogen-derived peptides, AE17 tumor-bearing mice received 100 µl of SQ vaccine containing 20-mer peptides listed in Supplementary Table 1 formulated with K3/c-di-AMP, while IT vaccines (30 µl) containing the same pathogen-derived peptides and K3/c-di-AMP were administered when tumor volume reached 50 mm^3^ size. Vaccinations using pathogen-derived peptides were performed according to the P/T neopeptide vaccination schedule above.

### Detection of neopeptide-specific T cells in spleens

Mice were vaccinated with the SQ AE17 neopeptide vaccine once a week for three weeks. Mice receiving SQ saline were used as a control. One week after the last SQ vaccination, spleens were harvested, mechanically dissociated, filtered with a 40-µm cell strainer and red blood cells were lysed using ACK lysis buffer (ammonium chloride (Sigma-Aldrich), potassium bicarbonate (Sigma-Aldrich) and EDTA (Sigma-Aldrich)). After washing, cells were counted. For intracellular cytokine staining of neopeptide-specific T cells, 2 × 10^6^ live cells were further restimulated with the AE17 neopeptides (4 µg/ml per peptide) at 37°C for 24 h, with GolgiPlug and GolgiStop (BD Biosciences) added during the last 6 h. Cells were then stained for intracellular cytokines and flow cytometry analysis as described below.

### Tumor tissue processing

For analysis of immune cells in tumors or tdLNs by flow cytometry and scRNA-seq, AE17 tumor-bearing mice received the SQ neopeptide vaccination once a week for three times. When tumor volume reached 100 mm³ size, mice received a single IT neopeptide vaccination. One, three, or seven days after the IT vaccination, mice were euthanized, and tumors and tdLNs were harvested. tdLNs were mechanically dissociated, filtered through a 40-µm cell strainer, and counted. Tumor tissues were weighed, minced into small fragments, and enzymatically digested using Tumor Dissociation Kit, mouse and gentleMACS Dissociator (Miltenyi Biotec) according to the manufacturer’s protocol. Digested tumor cells were washed with PRMI media and treated with red blood cell ACK lysis buffer. Cells were then filtered with a 40-µm cell strainer twice and counted using ReadyProbe Cell Viability Imaging Kit, Blue/Green (Invitrogen) following a 20-min incubation. Single-cell suspensions of tdLNs and digested tumors were either used freshly or cryopreserved in freezing medium (FBS with 10% DMSO) and stored at -80°C or liquid nitrogen until use. For intracellular cytokine staining of neopeptide-specific T cells in tumors and tdLNs, 1 × 10^6^ live cells of cryopreserved single-cell suspension were further restimulated with the AE17 neopeptides (5 µg/ml per peptide) in the presence of 0.5 × 10^5^ DC2.4 cells at 37°C for 24 h, with GolgiPlug and GolgiStop added during the last 6 h. Cells were then stained for intracellular cytokines and flow cytometry analysis as described below.

### Flow cytometry

For flow cytometric analysis, 2 × 10^6^ live cells from fresh tumor single-cell suspensions or restimulated cells from spleens, tdLNs, or digested tumors were washed with FACSWash buffer (HBSS (Gibco) containing 3% horse serum (Sigma-Aldrich), 0.02% sodium azide (Sigma-Aldrich), and 2.5mM calcium chloride (CaCl2, Thermo Fisher Scientific)) and blocked with anti-mouse CD16/CD32 antibody (BioLegend) for 10 min at 4°C. Cells were then stained with Annexin V (BD Biosciences) and the following surface antibodies listed in Supplementary Table 2 for 20 min at 4°C. For intracellular staining for cytokines, cells were washed with FACSWash buffer and fixed with IC Fixation Buffer (eBiosciences) containing 2.5mM CaCl2 for 60 min at 4°C. Cells were then washed with 1× permeabilization buffer and stained for cytokines using the antibodies listed in Supplementary Table 2 in 1× permeabilization buffer for 45 min at 4°C. After washing with 1× permeabilization buffer twice, cells were fixed with 1% paraformaldehyde (PFA, Merck) in PBS containing 2.5mM CaCl2. For transcription factor staining, cells were washed with FACSWash buffer and fixed and permeabilized using Foxp3/Transcription Factor Staining Buffer Set (eBiosciences) for 60 min at 4°C. Cells were then washed with 1× permeabilization buffer and stained for transcription factors using the antibodies listed in Supplementary Table 2 in 1× permeabilization buffer for 45 min at 4°C. After washing with 1× permeabilization buffer twice, cells were fixed with 1% PFA containing 2.5mM CaCl2. Samples of splenocytes were acquired on a BD LSRFortessa Cell Analyzer (BD Biosciences) and analyzed with Flowjo software v10.10.0 (BD Biosciences). Samples of digested tumors and tdLNs were acquired on a 5-laser Cytek Aurora spectral flow cytometer (Cytek Biosciences) and analyzed using OMIQ software (Dotmatics). Absolute cell numbers were calculated using flow cytometry data combined with the total number of live cells per gram of tumor tissue, determined by cell counting and tumor weight measurement.

### Immunofluorescence staining of tumors

For immunofluorescence staining, snap-frozen tumor tissues harvested one or seven days after a single IT vaccination following SQ priming were used. Frozen tumors were embedded with O.C.T. compounds (Sakura Finetek) and sectioned at 10 μm thickness using a Leica CM3050 S Cryostat (Leica Biosystems). Tumor sections were fixed in pre-chilled methanol at -20°C for 30 min. Following fixation, sections were blocked with blocking buffer containing of 3× SSC buffer, 2% BSA, and anti-mouse CD16/CD32 antibody for 15 min at room temperature. Sections were then stained with antibody solution containing 3× SSC buffer (Sigma-Aldrich), 2% Bovine Serum Albumin (BSA, Sigma-Aldrich), 4′,6-Diamidine-2′-phenylindole dihydrochloride (DAPI, Invitrogen) and the antibodies listed in Supplementary Table 2 for 1 h at room temperature. After washing with wash buffer (3× SSC buffer and 2% BSA), stained sections were mounted using ProLong™ Gold Antifade Mountant (Invitrogen). Images were acquired on a Zeiss AxioImager M2 microscpoe (ZEISS) and analyzed using ZEN Lite software (ZEISS).

### scRNA-seq sample preparation and sequencing

For scRNA-seq, frozen vials of tumor single-cell suspension harvested seven days post-IT vaccination were used. After thawing and washing, cells were blocked with anti-mouse CD16/CD32 antibody for 10 min at 4°C and stained with Apotracker Green (BioLegend) and the TotalSeq C antibodies (Supplementary Table 2) for 30 min at 4°C. Simultaneously, to multiplex samples some samples were also hashed using TotalSeq C hashtag antibodies from BioLegend (TotalSeq-C0301, 155861; TotalSeq-C0302, 155863; TotalSeq-C0303, 155865). Then, live cells (gated as Apotracker^-^) were sorted using a FACSAria III Cell Sorter (BD Biosciences) and subsequently subjected to sequencing. Hashed samples were pooled and cell suspensions were loaded into Chromium Next GEM Chip K (10X Genomics) for the 5’ capture. scRNA-seq libraries were then prepared using the Chromium Next GEM Single Cell 5’ Kit v2 (10X Genomics) and Chromium Single Cell Mouse TCR Amplification kit (10X Genomics) according to the manufacturer’s protocols. Then, the libraries were sequenced on a NovaSeq 6000 Sequencing System (Illumina).

### Data processing of scRNA- and TCR-seq

The raw sequencing data were processed simultaneously using the ‘cellranger multì pipeline with cellranger v7.2.0^50^ with mus musculus mm10-1.2.0 and vdj_GRCm38_alts_ensembl-2.2.0 as references. The count matrices were imported into R v4.4.2 and processed as a Seurat object with multiple assays: gene expression (RNA), antibody-derived tags (ADT; CD4, CD8a, CD11b, CD11c, CD19, F4-80, NK1.1), and hashtag oligos (HTO), using Seurat v5.2.1^51^. To separate multiplexed two samples in a library pool by their hashtags, the HTO assay was normalized with centered log-ratio (CLR) transformation across features and then demultiplexed using scDemultiplex v0.1.1^52^. The cells were categorized into Hashtag1, Hashtag2, Negative, and Doublet. Negative and Doublet cells were removed for true single cell resolution. As quality control, dying cells (‘percent.mt‘ > 25) and potential doublets (‘nFeature_RNÀ > 6000 or ‘nCount_RNÀ > 50000) were removed. The data was log-normalized and then scaled on the 2000 most variable genes, and principal component analysis (PCA) is performed. Data integration using the reciprocal PCA method (‘ RPCAIntegration‘) was performed with the first 20 PCs to correct the batch effects. Immune cells were separated based on their positive expression of *Ptprc* (encoding CD45). TCR clonotypes for a pair of TRA and TRB from the V(D)J library (‘airr_rearrangement.tsv‘) were added to the Seurat object using DALI^53^::Read10X_AIRR.

### Immune cell clustering and cell type identification

Immune cells with non-zero *Ptprc* gene expression were subject to clustering based on Euclidean distance of their seven ADT levels (CITE-seq) using Leiden algorithm with resolution 0.2. In ‘adt_snn_res.0.2‘, metaclusters 1 and 2 were positive for *Cd3e* and *Cd3d* therefore separated as T cell cluster, whereas the rest of metaclusters (0, 3, and 4) were separated as Myeloid cluster. Subsequently, the cell subsets were subject to clustering based on their RNA expression levels. They were separately normalized, scaled to the redefined 2000 variable features, SNN graph of the first 20 PCs were calculated, and clustered based on their RNA levels using Leiden algorithm with resolution 0.8. In T cell cluster, four subclusters were negative for *Cd3e* and *Cd3d*. Additionally these clusters were positive for fibroblast-related genes such as *Serpinh1*, NK cell-related genes such as *Klrb1c*, antigen-presenting cell-related genes such as *H2-Aa*, and mast cell-related genes such as *Cpa3*, respectively, suggesting that these cell populations are non-T cells. Therefore, these clusters are removed for further T cell analysis. The filtered T cell subset was re-clustered and manually annotated based on their expression levels of canonical makers and top 10 DEGs. In Myeloid cluster, two subclusters were positive for T cell-related genes such as *Cd3e* and fibroblast-related genes such as *Serpinh1*, respectively, therefore these clusters were removed for further myeloid cell analysis. The filtered myeloid cell subset was re-clustered and manually annotated based on their expression levels of canonical makers and the top 10 DEGs.

### CDR3 amino acid sequence clustering

T cell cluster with clonotype information were re-clustered on their beta chain’s Complementarity Determining Region 3 (CDR3) amino acid sequence (‘vdj.cdr3‘) using clusTCR^54^ with the following option values: distance_metric = “hamming”, method = “two-step”. The top 30 CDR3 motifs were chosen based on the frequency in all cells that participated in the clustering process regardless of their cell types. For visualization, the original cell types were merged into a coarser resolution. CD8_TEX-term, CD8_TEX-prolif, and CD8_TPEX were merged and referred to as exhausted CD8^+^ T cell subsets. CD4_Th1, CD4_Th1_CD40L+, CD4_Th1_OX40+, CD4_Th1_ISG+, and CD4_Th1_Ki67+ were merged and referred to as Th1 subsets. Gini index was calculated in each cell type using the amino acid sequence within each cell type using DALI::ClonotypeDiversity with the following options: algorithm = c(“gini”), group.by = “orig.ident”, by.group = TRUE, use.sequence = TRUE, sequence.type = “AA”, chain = “VDJ”.

### Differentially expression analysis and GO analysis

For differential gene expression analysis (DEA) on Myeloid cluster, DEA was performed per cell type on their normalized counts between two conditions using Seurat implementation of MAST (Model-based Analysis of Single-cell Transcriptomics)^55^, and the cellular detection rate (CDR), the fraction of genes expressed in a cell, has been corrected (‘latent.var = CDR‘). For GO analysis enriched in the Myeloid cluster, the upregulated top-50 DEGs in the P/T vaccination group in comparison with the SQ saline + IT saline group based on pseudo bulk data set were imported into the web-based MetaScape portal ^56^.

### Real Time-qPCR of tumor tissues

To examine gene expression in tumor tissues by qPCR, frozen digested tumors or frozen tumor tissues harvested one or seven days post-IT vaccination following SQ priming were used. Frozen tumor tissues embedded with O.C.T. compounds were sectioned using a Leica CM3050 S Cryostat. RNA was isolated from tumor sections or 1 × 10^6^ cells derived from digested tumors using TRIzol Reagent (Invitrogen) according to the manufacturer’s protocol. Subsequently, cDNA was synthesized using High-Capacity cDNA Reverse Transcription Kit (Applied Biosystems) according to the manufacturer’s protocol. RT-qPCR was then performed on a QuantStudio 5 Real-Time PCR System (Applied Biosystems) using the following Taqman Gene Expression assays: Gapdh (Mm99999915), Cxcl9 (Mm00434946), Cxcl10 (Mm00445235), Ifnb1 (Mm00439552), and Ifng (Mm01168134). Fold change of the gene expression was calculated using the 2^(-ΔΔCt) method. Briefly, ΔCt values were calculated by subtracting the average of Ct of an internal housekeeping gene (*Gapdh*; technical triplicates) from the average Ct of the target gene for each sample (technical triplicates per sample). ΔΔCt values were then calculated by subtracting the ΔCt of the SQ saline + IT saline group from the ΔCt of each treatment group. Fold changes were then calculated as 2^(-ΔΔCt), and log₂ fold change was subsequently calculated as log₂(2^(-ΔΔCt)).

### Statistical analysis

Data were first evaluated for normal distribution by Shapiro-Wilk normality test. If data was normally distributed, ordinary one-way ANOVA with Tukey’s multiple comparisons test or two-tailed unpaired Student’s t-test were used. If not normally distributed, Kruskal-Wallis test with Dunn’s multiple comparisons test or two-tailed unpaired Mann-Whitney U-test were used. Fisher’s exact test was used to compare the frequencies of tumor ulcers across the groups. For statistical analysis of average tumor growth and Kaplan-Meier survival curves, two-way ANOVA with Bonferroni multiple comparisons test and log-rank (Mantel-Cox) test, respectively, were used. The number of samples (n) per group and independent experiments and statistical tests used are stated in the figure legends. Statistical analyses were performed using GraphPad Prism (v 9.0.0). *P* values < 0.05 were considered significant.

## ACKNOWLEDGEMENTS

This work was supported by Department of Immunology of Erasmus University Medical Center and the Dutch Cancer Society (KWF grant 12837) and in part by IMSUT International Joint Usage/Research Center Project (K22-3063 and K25-3190) from The Institute of Medical Science, The University of Tokyo (IMSUT).

## AUTHOR CONTRIBUTIONS

K.H. and P.D.K. conceptualized the study, designed the experiments, and interpreted the results. C.S., Y.M., B.T. and K.J.I. contributed to data analysis and interpretation. K.H., M.W., I.B.-H., and M.v.M. conducted experiments and analyzed data. K.H., M.W., and E.B. performed scRNA-seq. Y.L., K.H., and H.J.G.v.d.W. analyzed the scRNA-seq data. K.H., C.S., and P.D.K. wrote and revised the manuscript. All authors read, revised and approved the manuscript.

---

**Extended Data Fig. 1.**
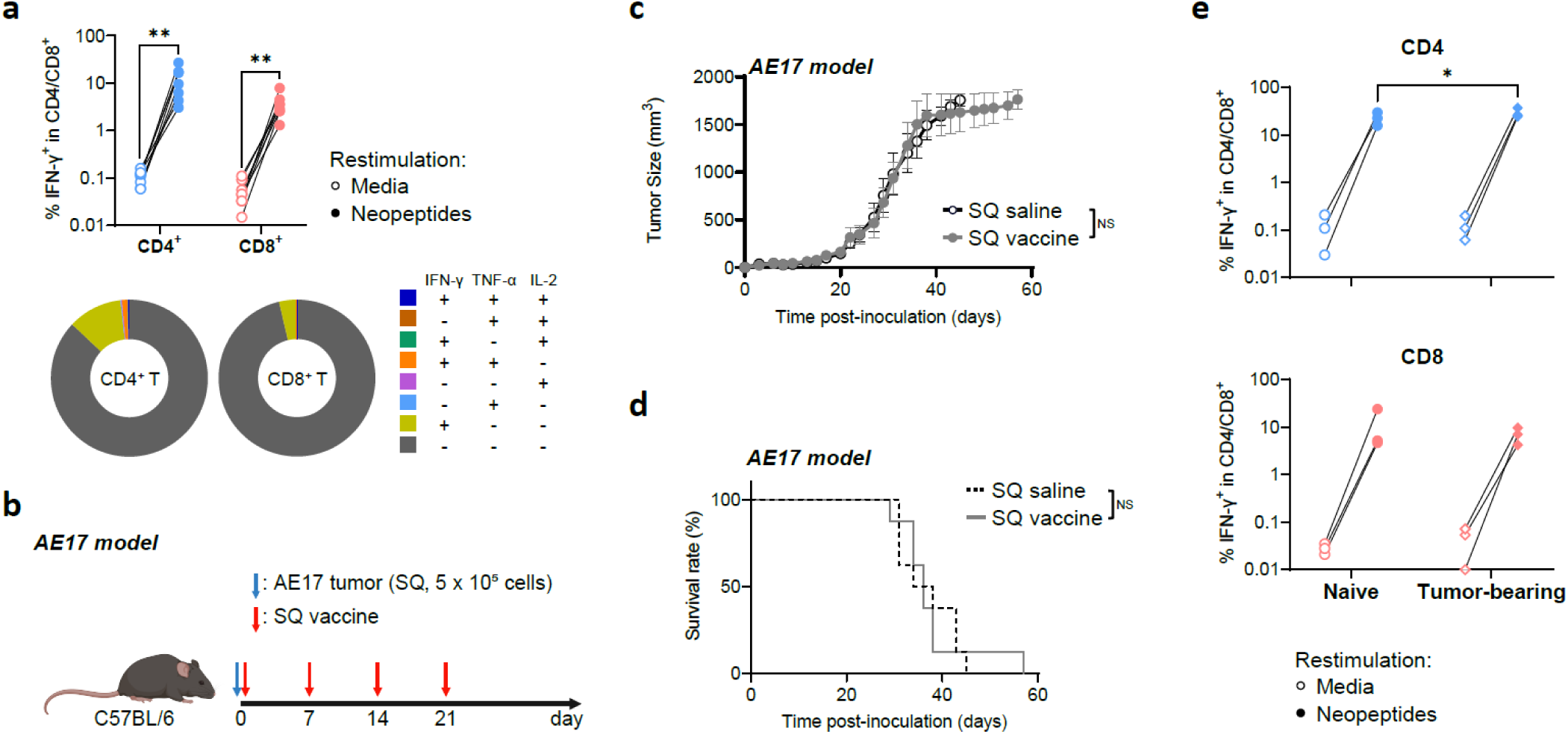
SQ neopeptide vaccination alone fails to control tumor growth in the AE17 tumor model, **a,** Top, frequencies of neopeptide-specific CD4^+^ or CD8^+^ T cells in spleens of SQ-vaccinated mice. Neopeptide-specific cytokine production was assessed upon restimulation with the corresponding neopeptides; media condition served as a negative control. Bottom, distribution of cytokine-producing (IFN-γ, TNF-α, and/or IL-2^+^) cells within CD4^+^ or CD8^+^ T cells in the SQ. vaccination group (n = 8, average), b, Experimental design to examine the efficacy of SQ vaccination in the AE17 tumor model. C57BL/6J mice were inoculated with AE17 tumor cells (SQ, 0.5 x 10^6^ cells) on day 0. SQ neopeptide vaccines were administered **on** days 0, 7, 14, and 21. **c, d,** Average tumor growth curve **(c)** and Kaplan-Meier survival curve **(d)** of mice bearing AE17 tumors treated as indicated, e, Frequencies of neopeptide-specific CD4^+^ (top) or CD8^+^ (bottom) T cells in spleens of SQ-vaccinated mice (Naïve) or SQ-vaccinated tumor-bearing mice (Tumor­bearing). Neopeptide-specific cytokine production was assessed upon restimulation with the corresponding neopeptides; media condition served as a negative control. Data in **a** are from one experiment (n = 8 mice per group) and were statistically analyzed by two-tailed unpaired Student’s t-test for each CD4^+^ and CD8^+^T cells. Average tumor growth curve was statistically analyzed by two-way ANOVA with Bonferroni multiple comparisons test as of the end point of the SQ saline + IT saline group **(c).** Survival curve was statistically analyzed by log-rank (Mantel-Cox) test **(d).** Data in e are from one experiment (n = 3 mice per condition) were statistically analyzed by two-way ANOVA with Bonferroni multiple comparisons test, *p < 0.05; NS, not significant.

**Extended Data Fig. 2.**
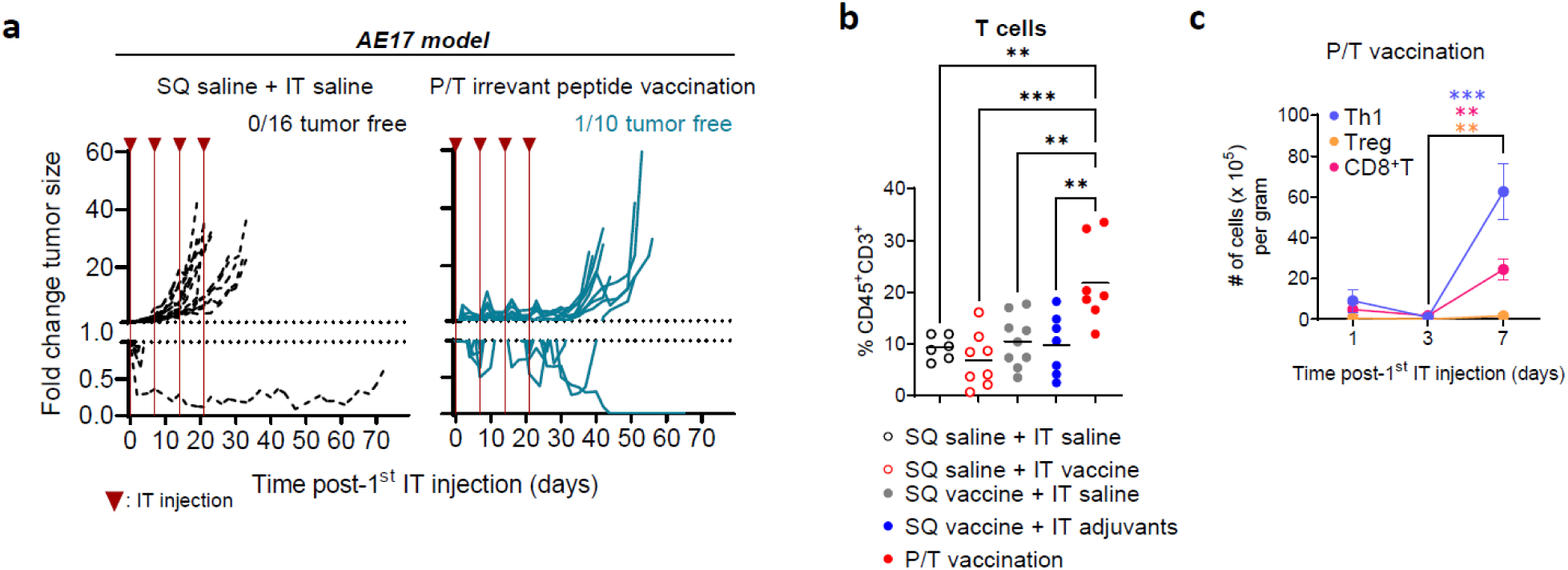
P/T vaccination using irrelevant pathogen-derived peptides does not achieve prolonged tumor control, **a,** Fold change in tumor size relative to the tumor size at the time of the 1^st^ IT injection in the SQ saline + IT saline and P/T viral vaccination groups in the AE17 tumor model. The data shown represent the time course post-1^st^ IT injection. Inverted triangles indicate IT injection time points, **b,** Frequency of tumor-infiltrating T cells, defined as CD45^+^CD3^+^ among live singlets, per group at day 7 after a single IT vaccination and after SQ priming. Data from the SQ saline + IT saline and P/T vaccination groups are also shown in Fig. 2b. **c,** Kinetics of Thl, Treg, and CD8^+^ T cells within tumors harvested on days 1, 3, or 7 after a single IT vaccination in the P/T vaccination group (mice were SQ primed), shown as the absolute number per gram of tumor tissue, based on flow cytometric analysis shown in Fig. 2g (n = 3-8 per time point from two independent experiments). Data at the day 7 time point are also shown in Fig. 2g.

**Extended Data Fig. 3.**
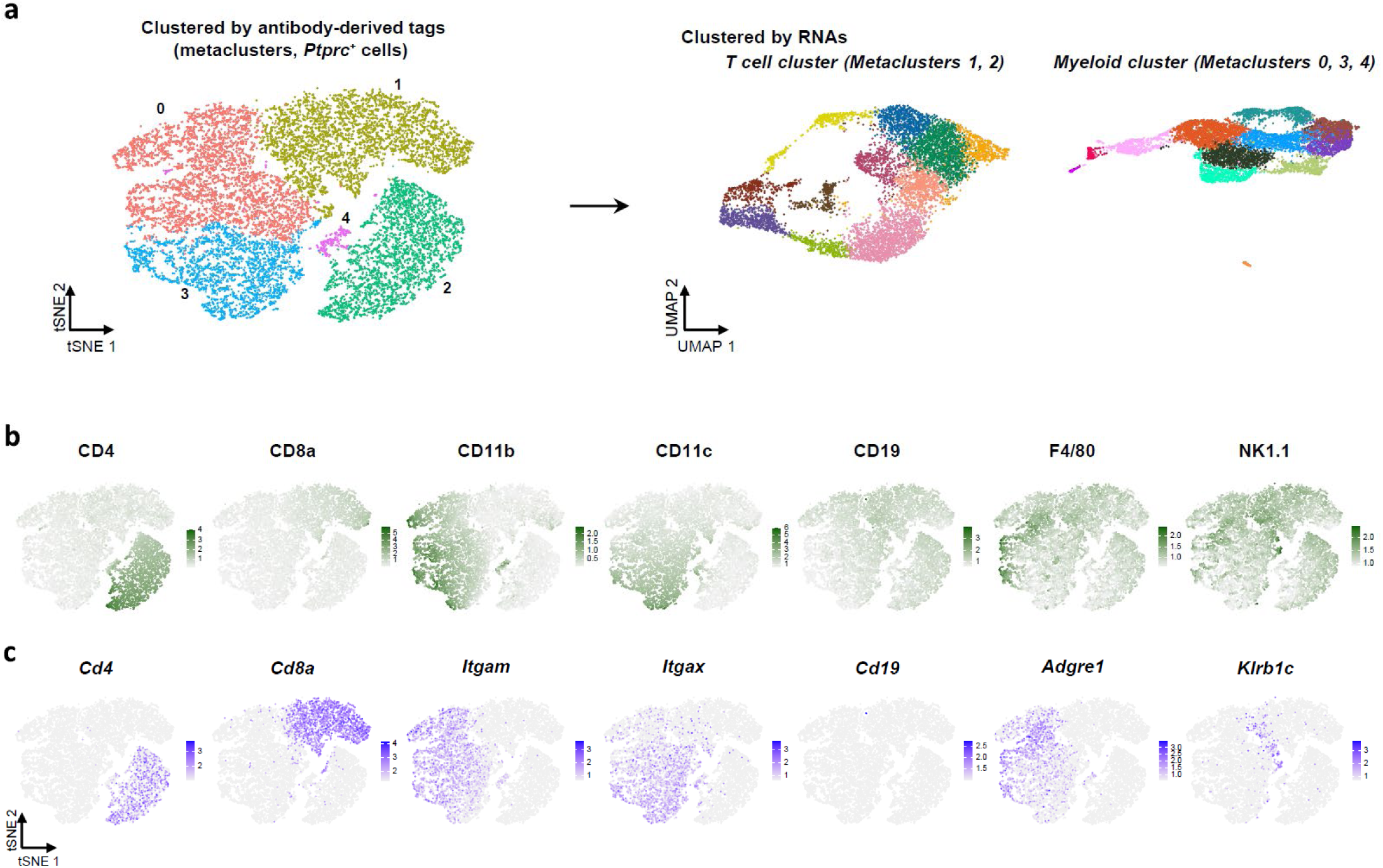
Identification of the T cell and myeloid clusters in scRNA-seq of tumors, **a,** tSNE plot showing metaclustering of *Ptprc^+^* (CD45^+^) immune cells based on ADT protein expressions. Metaclusters 1 and 2 were assigned as the T cell cluster, while metaclusters 0, 3, and 4 were assigned as the myeloid cluster, **b,** tSNE plots of immune cells from **a,** showing normalized expression of proteins based on ADT. **c,** tSNE plots of immune cells from **a,** showing normalized expression of genes corresponding to the proteins **in b.**

**Extended Data Fig. 4.**
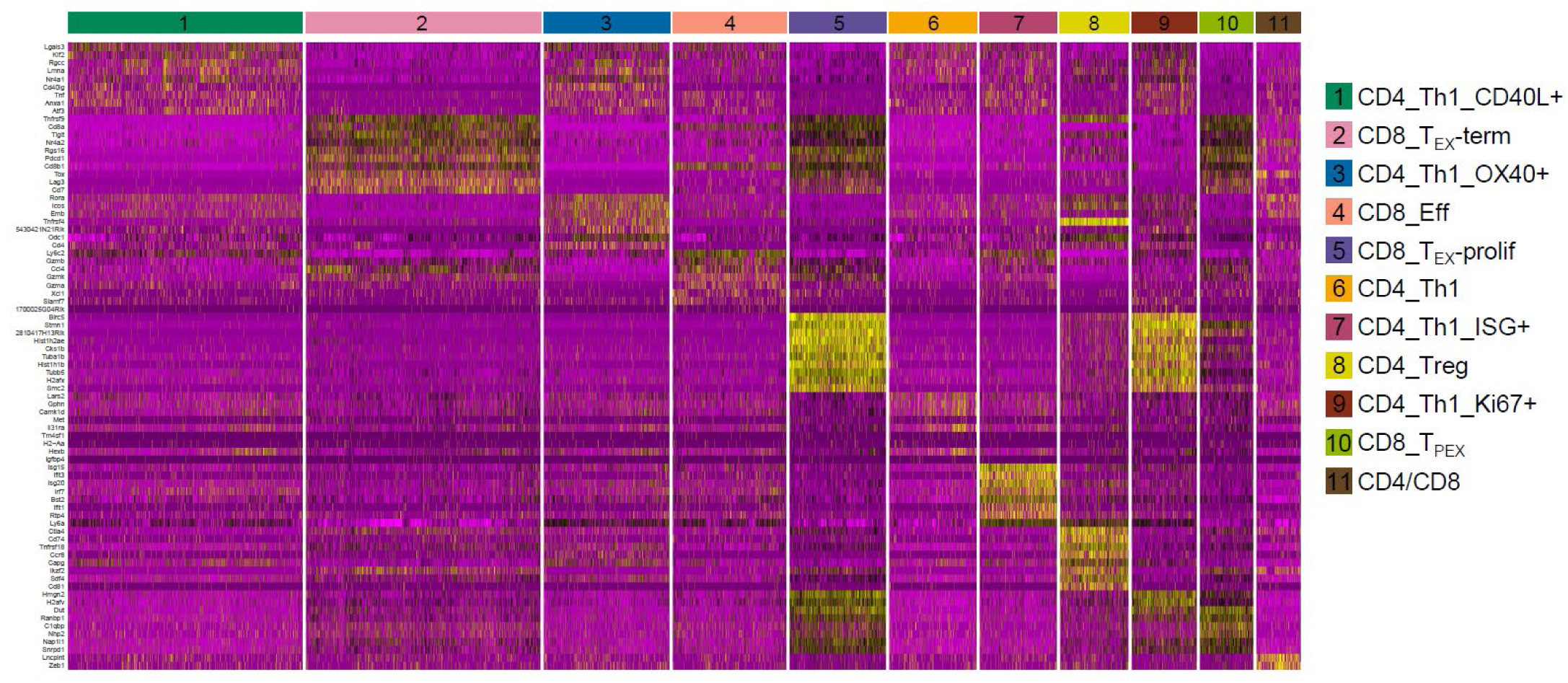
Characterization of T subsets in the T cell cluster of scRNA-seq. Heatmap showing the top 10DEGs in each T cell subcluster.

**Extended Data Fig. 5.**
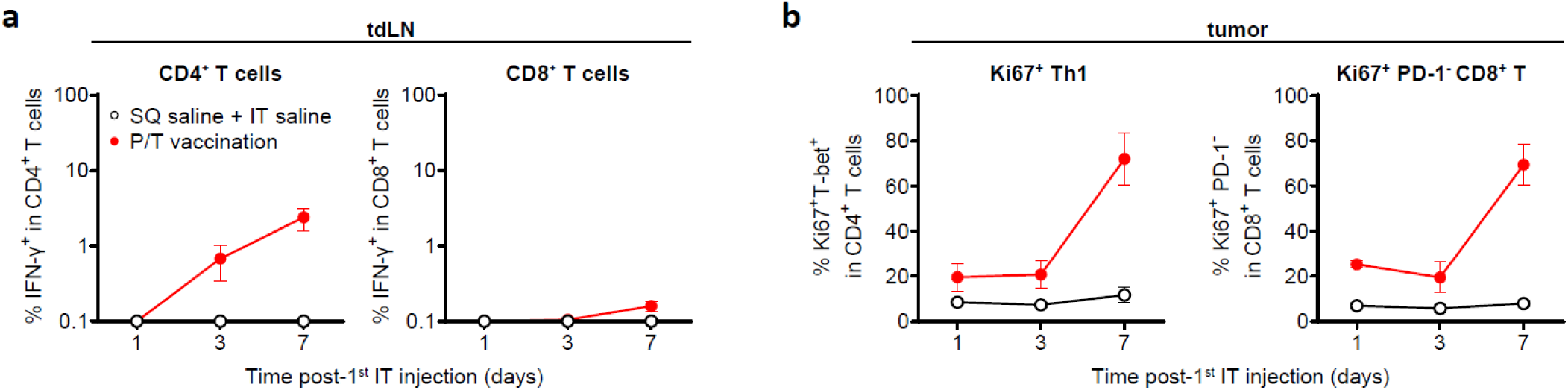
Proliferating T cells accumulate in tumors at day 7 after a single IT vaccination in the P/T vaccination group, **a**, Kinetics of neopeptide-specific CD4^+^ (left) or CD8^+^ (right) T cells, shown as frequencies within CD4^+^ or CD8^+^ T cells in tdLNs harvested on days 1, 3, or 7 days after a single IT vaccination in the SQ saline + IT saline and P/T vaccination groups. Neopeptide-specific IFN-y production was assessed upon restimulation with the corresponding neopeptides, b, Kinetics of Ki67^+^Thl cells (left) or Ki67^+^ PD-1 CD8^+^ T cells (right), shown as frequencies within CD4^+^ or CD8^+^ T cells in tumors harvested on days 1, 3, or 7 days after a single IT vaccination in the SQ saline + IT saline and P/T vaccination groups. Cells were analyzed by flow cytometry.

**Extended Data Fig. 6.**
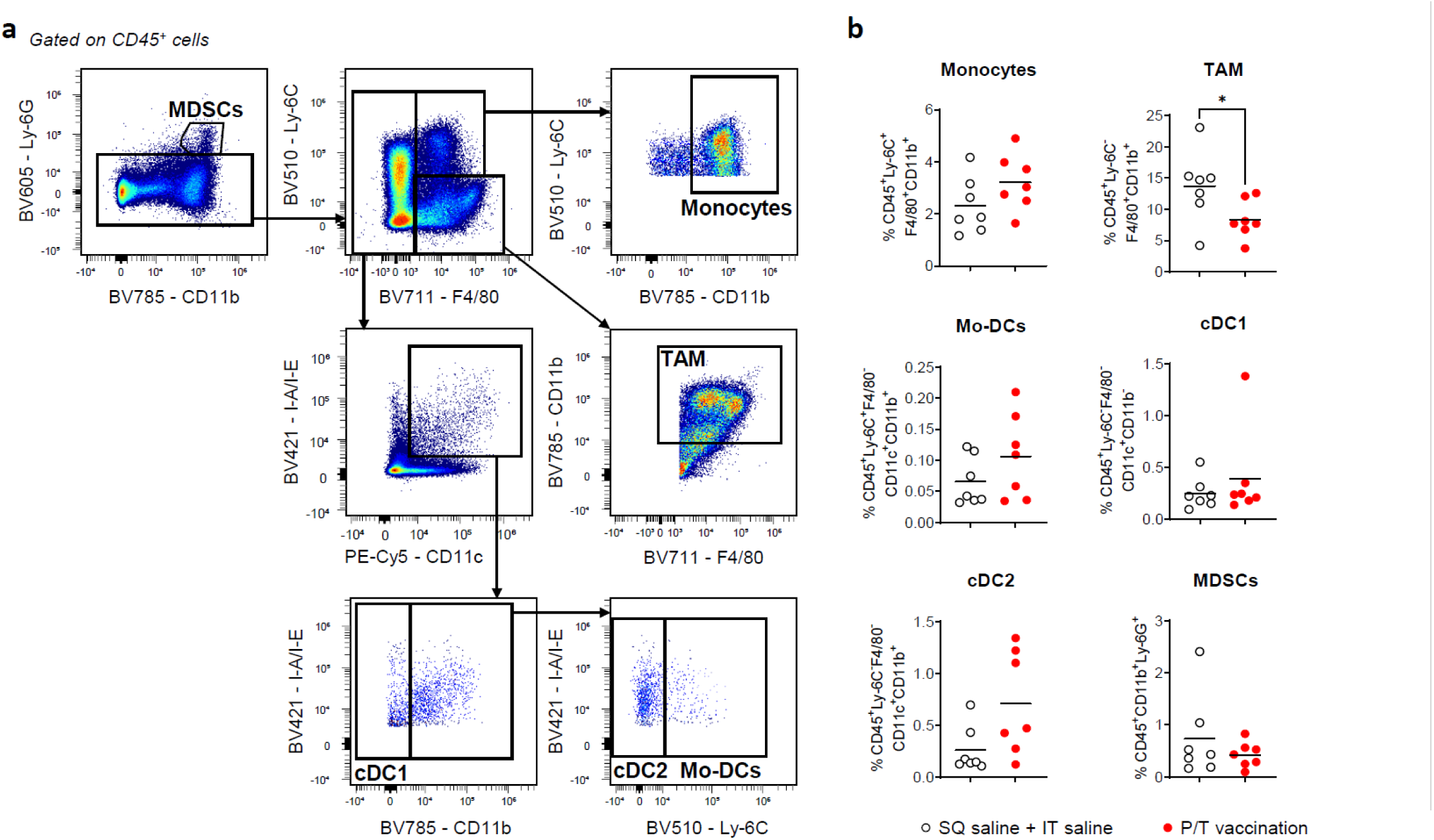
P/T neopeptide vaccination reduces TAMs within the TME. **a**, Gating strategy on CD45+ cellsfor characterization of myeloid cell subsets by flow cytometry. MDSCs, myeloid-derived suppressor cells; TAM, tumorassociated macrophages; cDC1, conventional dendritic cells (DC) 1, cDC2, conventional DC 2, Mo-DCs, monocytederived DCs. **b**, Frequencies of the indicated myeloid cell subsets per group at day 7 after a single IT vaccination in theSQ saline + IT saline and P/T vaccination groups. Horizontal lines in b represent the mean. Data were statisticallyanalyzed by two-tailed unpaired Student’s t-test or two-tailed unpaired Mann-Whitney U-test. \**p* < 0.05.

**Extended Data Fig. 7.**
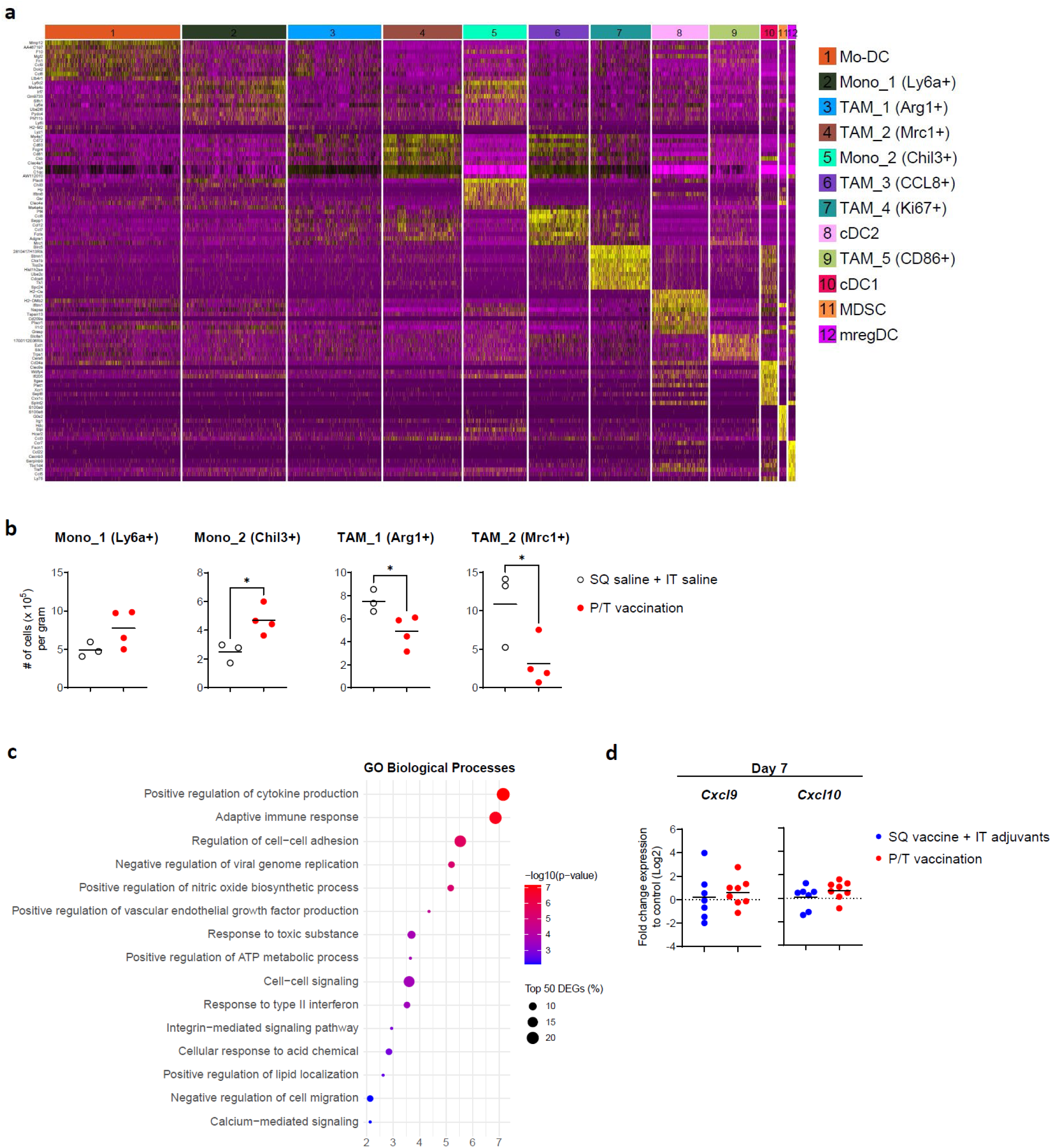
P/T neopeptide vaccination reduces dominant TAM subsets within the TME. **a,** Heatmap of scRNASeq showing the top 10 DEGs in each myeloid subcluster, **b,** Absolute numbers of Mono_l (Ly6a+), Mono_2 (Chil3÷), TAM_l (Argl÷), and TAM_2 (Mrcl+) per gram of tumor tissue at 7 days after a single IT vaccination in the SQ saline + IT saline and P/T vaccination groups, based on scRNA-seq of the myeloid cluster shown in Fig. 5a. c, Analysis of gene ontology (GO) biological processes comparing the myeloid cluster from the SQ saline + IT saline versus P/T vaccination groups based on top 50 DEGs in pseudo bulk data set. d, Fold change in *Cxcl9* (left) and *CxcIlO* (right) expression, analyzed by qPCR, in tumors harvested at day 7 after a single IT vaccination in the SQ vaccine + IT adjuvants and P/T vaccination groups (n= 7 or 8 per group from three independent experiments). Fold change was normalized to the SQ saline + IT saline group and converted to Log2 scale. Horizontal linesin **b** and **d** represent the mean. Data were statistically analyzed two-tailed unpaired Student’s t-test. *\*p* < 0.05.

**Table S1.**
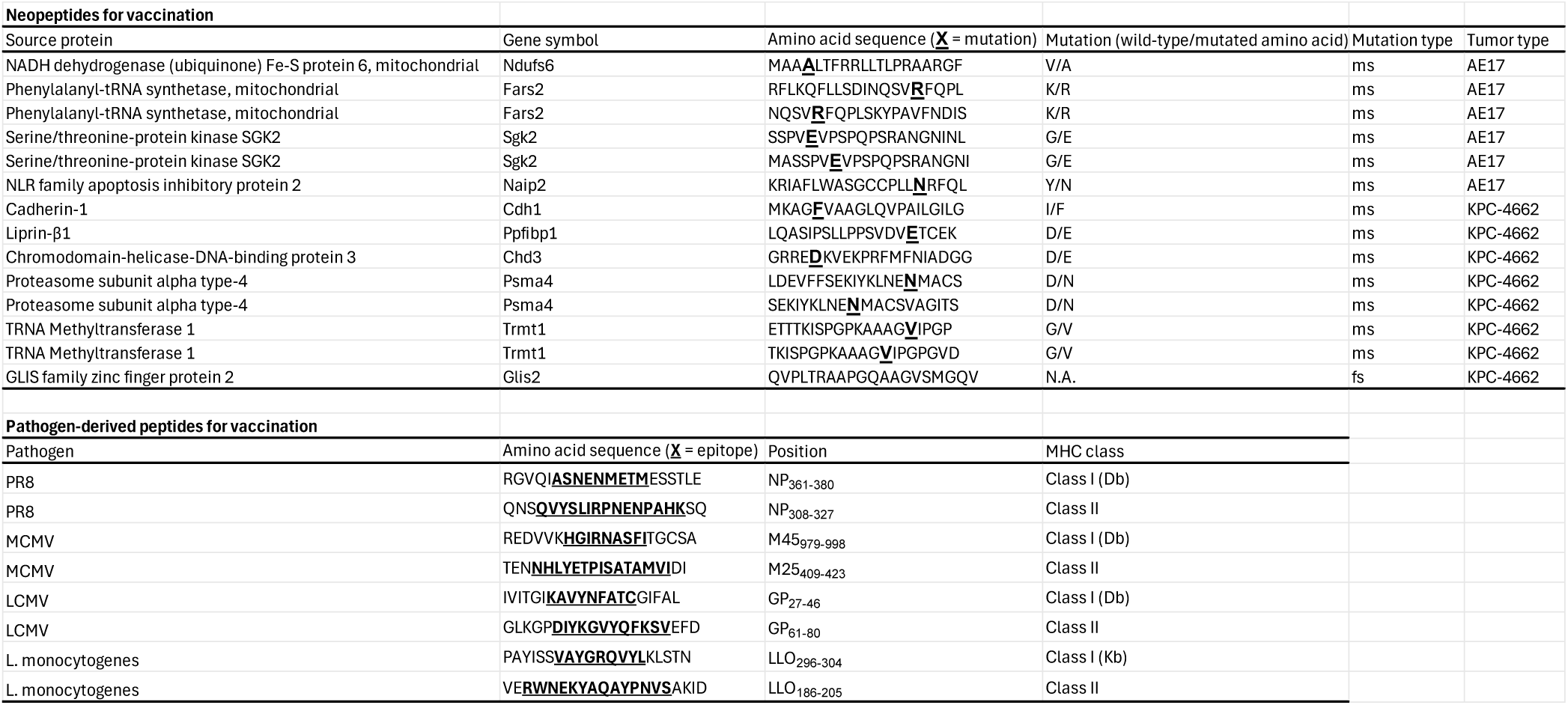
Neopeptides and pathogen-derived peptides used for vaccination. For neopeptides, the table lists source proteins, gene symbols, amino acid sequences containing mutations, mutation types (ms, missense; fs, frameshift), and tumor types. For pathogen-derived peptides, the table list pathogens, epitope-containing amino acid sequences, epitope positions, and MHC class restriction. N.A., not applicable; PR8, Influenza A virus (A/Puerto Rico/8/1934, H1N1); MCMV, murine cytomegalovirus; LCMV, lymphocytic choriomeningitisvirus; L. monocytogenes, Listeria monocytogenes.

**Table S2.**
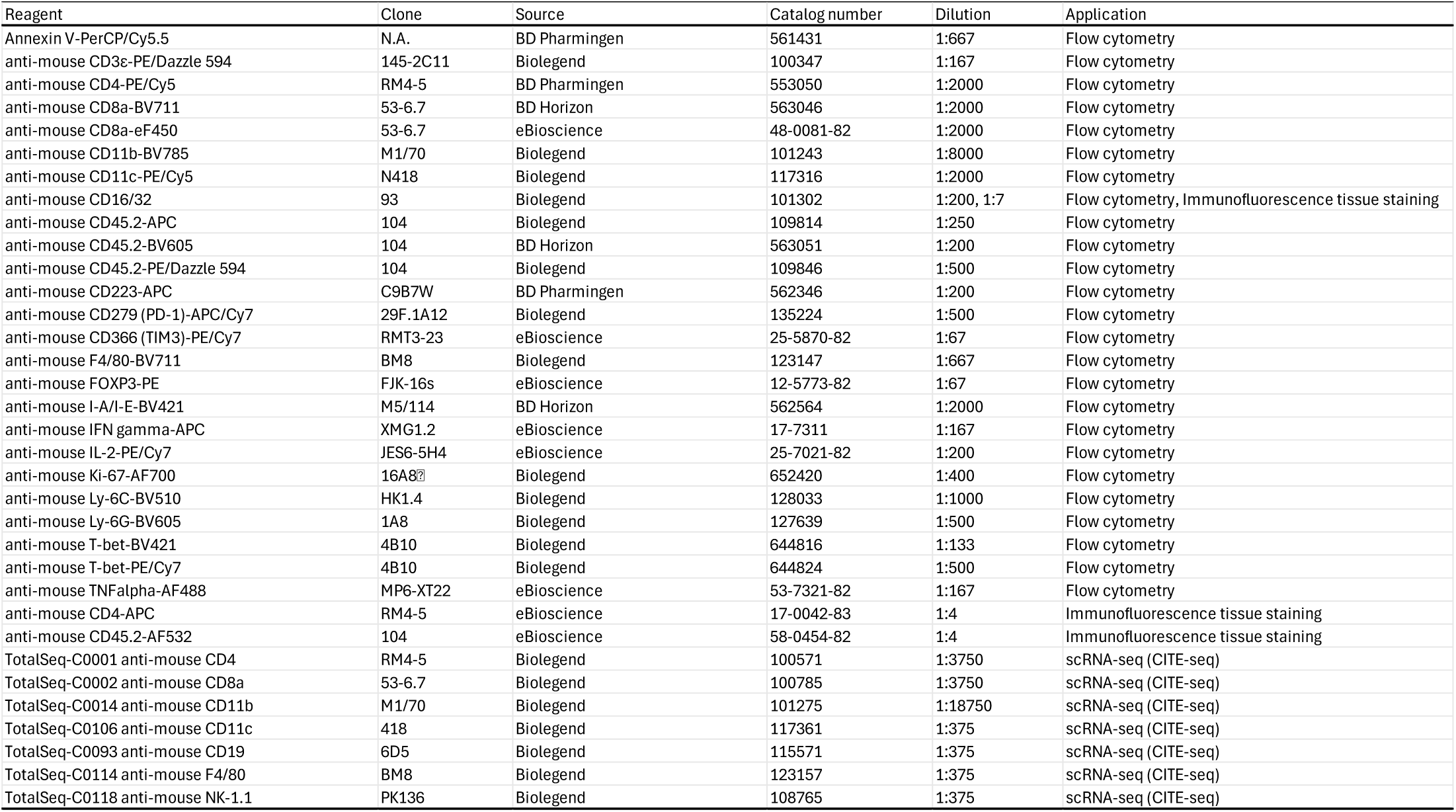
Annexin V and antibodies used for flow cytometry, immunofluorescence tissue staining, and CITE-seq for scRNA-seq. The table lists Annexin V and antibodies, including their clones, sources, catalog numbers, and dilution factors used for flow cytometry, immunofluorescence tissue staining, and CITE-seq for scRNA-seq. N.A., not applicable.

## REFERENCES

1 Katsikis, P. D., Ishii, K. J. & Schliehe, C. Challenges in developing personalized neoantigen cancer vaccines. Nat Rev Immunol 24, 213–227 (2024). 10.1038/s41577-023-00937-y

2 Hanahan, D., Michielin, O. & Pittet, M. J. Convergent inducers and effectors of T cell paralysis in the tumour microenvironment. Nat Rev Cancer (2024). https://doi.org/10.1038/s41568-024-00761-z [pii] 10.1038/s41568-024-00761-z

3 Wu, B., Zhang, B., Li, B., Wu, H. & Jiang, M. Cold and hot tumors: from molecular mechanisms to targeted therapy. Signal Transduct Target Ther 9, 274 (2024). https://doi.org/10.1038/s41392-024-01979-x [pii] 1979 [pii] 10.1038/s41392-024-01979-x

4 Bonaventura, P. et al. Cold Tumors: A Therapeutic Challenge for Immunotherapy. Front Immunol 10, 168 (2019).

5 Franken, A. et al. CD4(+) T cell activation distinguishes response to anti-PD-L1+anti-CTLA4 therapy from anti-PD-L1 monotherapy. Immunity (2024). https://doi.org/S1074-7613(24)00083-9 [pii] 10.1016/j.immuni.2024.02.007

6 Borst, J., Ahrends, T., Babala, N., Melief, C. J. M. & Kastenmuller, W. CD4(+) T cell help in cancer immunology and immunotherapy. Nat Rev Immunol 18, 635–647 (2018). https://doi.org/10.1038/s41577-018-0044-0 [pii] 10.1038/s41577-018-0044-0

7 Wei, S. C. et al. Distinct Cellular Mechanisms Underlie Anti-CTLA-4 and Anti-PD-1 Checkpoint Blockade. Cell 170, 1120–1133 e1117 (2017). https://doi.org/S0092-8674(17)30831-0 [pii] 10.1016/j.cell.2017.07.024

8 Saxena, M., van der Burg, S. H., Melief, C. J. M. & Bhardwaj, N. Therapeutic cancer vaccines. Nat Rev Cancer 21, 360–378 (2021). 10.1038/s41568-021-00346-0

9 Hegde, P. S. & Chen, D. S. Top 10 Challenges in Cancer Immunotherapy. Immunity 52, 17–35 (2020). https://doi.org/S1074-7613(19)30530-8 [pii] 10.1016/j.immuni.2019.12.011

10 Burn, O. K., Prasit, K. K. & Hermans, I. F. Modulating the Tumour Microenvironment by Intratumoural Injection of Pattern Recognition Receptor Agonists. Cancers (Basel) 12 (2020). 10.3390/cancers12123824

11 Lin, M. J. et al. Cancer vaccines: the next immunotherapy frontier. Nat Cancer 3, 911–926 (2022). 10.1038/s43018-022-00418-6

12 Hong, W. X. et al. Intratumoral Immunotherapy for Early-stage Solid Tumors. Clin Cancer Res 26, 3091–3099 (2020). https://doi.org/1078-0432.CCR-19-3642 [pii] 10.1158/1078-0432.CCR-19-3642

13 Morisaki, T. et al. Lymph Nodes as Anti-Tumor Immunotherapeutic Tools: Intranodal-Tumor-Specific Antigen-Pulsed Dendritic Cell Vaccine Immunotherapy. Cancers (Basel) 14 (2022). https://doi.org/cancers14102438 [pii] cancers-14-02438 [pii] 10.3390/cancers14102438

14 Sahin, U. et al. Personalized RNA mutanome vaccines mobilize poly-specific therapeutic immunity against cancer. Nature 547, 222–226 (2017). 10.1038/nature23003

15 Kreiter, S. et al. FLT3 Ligand as a Molecular Adjuvant for Naked RNA Vaccines. Methods Mol Biol 1428163–175 (2016). 10.1007/978-1-4939-3625-0_11

16 Kreiter, S. et al. Intranodal vaccination with naked antigen-encoding RNA elicits potent prophylactic and therapeutic antitumoral immunity. Cancer Res 70, 9031–9040 (2010). https://doi.org/0008-5472.CAN-10-0699 [pii] 10.1158/0008-5472.CAN-10-0699

17 Tanyi, J. L. et al. Personalized cancer vaccine effectively mobilizes antitumor T cell immunity in ovarian cancer. Sci Transl Med 10 (2018). https://doi.org/10/436/eaao5931 [pii] 10.1126/scitranslmed.aao5931

18 Castro Eiro, M. D. et al. TLR9 plus STING Agonist Adjuvant Combination Induces Potent Neopeptide T Cell Immunity and Improves Immune Checkpoint Blockade Efficacy in a Tumor Model. J Immunol 212, 455–465 (2024). 10.4049/jimmunol.2300038

19 Dolfi, D. V. et al. Dendritic cells and CD28 costimulation are required to sustain virus-specific CD8+ T cell responses during the effector phase in vivo. J Immunol 186, 4599–4608 (2011). https://doi.org/jimmunol.1001972 [pii] 10.4049/jimmunol.1001972

20 Prokhnevska, N. et al. CD8(+) T cell activation in cancer comprises an initial activation phase in lymph nodes followed by effector differentiation within the tumor. Immunity (2022). https://doi.org/S1074-7613(22)00606-9 [pii] 10.1016/j.immuni.2022.12.002

21 Kyi, C. et al. Therapeutic Immune Modulation against Solid Cancers with Intratumoral Poly-ICLC: A Pilot Trial. Clin Cancer Res 24, 4937–4948 (2018). https://doi.org/1078-0432.CCR-17-1866 [pii] 10.1158/1078-0432.CCR-17-1866

22 Salazar, A. M., Erlich, R. B., Mark, A., Bhardwaj, N. & Herberman, R. B. Therapeutic in situ autovaccination against solid cancers with intratumoral poly-ICLC: case report, hypothesis, and clinical trial. Cancer Immunol Res 2, 720–724 (2014). https://doi.org/2326-6066.CIR-14-0024 [pii] 10.1158/2326-6066.CIR-14-0024

23 Nishii, N. et al. Systemic administration of a TLR7 agonist attenuates regulatory T cells by dendritic cell modification and overcomes resistance to PD-L1 blockade therapy. Oncotarget 9, 13301–13312 (2018). https://doi.org/24327 [pii] 10.18632/oncotarget.24327

24 Espinosa-Carrasco, G. et al. Intratumoral immune triads are required for immunotherapy-mediated elimination of solid tumors. Cancer Cell (2024). https://doi.org/S1535-6108(24)00193-4 [pii] 10.1016/j.ccell.2024.05.025

25 Giles, J. R., Globig, A. M., Kaech, S. M. & Wherry, E. J. CD8(+) T cells in the cancer-immunity cycle. Immunity 56, 2231–2253 (2023). https://doi.org/S1074-7613(23)00410-7 [pii] 10.1016/j.immuni.2023.09.005

26 Wei, C. et al. Tumor-associated macrophage clusters linked to immunotherapy in a pan-cancer census. NPJ Precis Oncol 8, 176 (2024). https://doi.org/10.1038/s41698-024-00660-4 [pii] 660 [pii] 10.1038/s41698-024-00660-4

27 Guimaraes, G. R. et al. Single-cell resolution characterization of myeloid-derived cell states with implication in cancer outcome. Nat Commun 15, 5694 (2024). https://doi.org/10.1038/s41467-024-49916-4 [pii] 49916 [pii] 10.1038/s41467-024-49916-4

28 Cheng, S. et al. A pan-cancer single-cell transcriptional atlas of tumor infiltrating myeloid cells. Cell 184, 792–809 e723 (2021). https://doi.org/S0092-8674(21)00010-6 [pii] 10.1016/j.cell.2021.01.010

29 Roychoudhuri, R., Eil, R. L. & Restifo, N. P. The interplay of effector and regulatory T cells in cancer. Curr Opin Immunol 33, 101–111 (2015). https://doi.org/S0952-7915(15)00032-1 [pii] 10.1016/j.coi.2015.02.003

30 Iglesias-Escudero, M., Arias-Gonzalez, N. & Martinez-Caceres, E. Regulatory cells and the effect of cancer immunotherapy. Mol Cancer 22, 26 (2023). 10.1186/s12943-023-01714-0

31 Kohli, K., Pillarisetty, V. G. & Kim, T. S. Key chemokines direct migration of immune cells in solid tumors. Cancer Gene Ther 29, 10–21 (2022).

32 Pan, M. et al. Targeting CXCL9/10/11-CXCR3 axis: an important component of tumor-promoting and antitumor immunity. Clin Transl Oncol 25, 2306–2320 (2023). https://doi.org/10.1007/s12094-023-03126-4 [pii] 10.1007/s12094-023-03126-4

33 Metzemaekers, M., Vanheule, V., Janssens, R., Struyf, S. & Proost, P. Overview of the Mechanisms that May Contribute to the Non-Redundant Activities of Interferon-Inducible CXC Chemokine Receptor 3 Ligands. Front Immunol 8, 1970 (2017). 10.3389/fimmu.2017.01970

34 Gubin, M. M. et al. High-Dimensional Analysis Delineates Myeloid and Lymphoid Compartment Remodeling during Successful Immune-Checkpoint Cancer Therapy. Cell 175, 1014–1030 e1019 (2018). https://doi.org/S0092-8674(18)31242-X [pii] 10.1016/j.cell.2018.09.030

35 Zitvogel, L., Galluzzi, L., Kepp, O., Smyth, M. J. & Kroemer, G. Type I interferons in anticancer immunity. Nat Rev Immunol 15, 405–414 (2015). https://doi.org/nri3845 [pii] 10.1038/nri3845

36 Kwart, D. et al. Cancer cell-derived type I interferons instruct tumor monocyte polarization. Cell Rep 41, 111769 (2022). https://doi.org/S2211-1247(22)01652-7 [pii] 10.1016/j.celrep.2022.111769

37 Temizoz, B. & Ishii, K. J. Type I and II interferons toward ideal vaccine and immunotherapy. Expert Rev Vaccines 20, 527–544 (2021). 10.1080/14760584.2021.1927724

38 Jneid, B. et al. Selective STING stimulation in dendritic cells primes antitumor T cell responses. Sci Immunol 8, eabn6612 (2023). 10.1126/sciimmunol.abn6612

39 Alspach, E. et al. MHC-II neoantigens shape tumour immunity and response to immunotherapy. Nature 574, 696–701 (2019). https://doi.org/10.1038/s41586-019-1671-8 10.1038/s41586-019-1671-8 [pii]

40 Cardenas, M. A. et al. Differentiation fate of a stem-like CD4 T cell controls immunity to cancer. Nature (2024). https://doi.org/10.1038/s41586-024-08076-7 [pii] 10.1038/s41586-024-08076-7

41 Kreiter, S. et al. Mutant MHC class II epitopes drive therapeutic immune responses to cancer. Nature 520, 692–696 (2015). https://doi.org/nature14426 [pii] 10.1038/nature14426

42 Bawden, E. G. et al. CD4(+) T cell immunity against cutaneous melanoma encompasses multifaceted MHC II-dependent responses. Sci Immunol 9, eadi9517 (2024). 10.1126/sciimmunol.adi9517

43 Kruse, B. et al. CD4(+) T cell-induced inflammatory cell death controls immune-evasive tumours. Nature 618, 1033–1040 (2023). https://doi.org/10.1038/s41586-023-06199-x [pii] 6199 [pii] 10.1038/s41586-023-06199-x

44 Montauti, E., Oh, D. Y. & Fong, L. CD4(+) T cells in antitumor immunity. Trends Cancer (2024). https://doi.org/S2405-8033(24)00157-2 [pii] 10.1016/j.trecan.2024.07.009

45 Brightman, S. E. et al. Neoantigen-specific stem cell memory-like CD4(+) T cells mediate CD8(+) T cell-dependent immunotherapy of MHC class II-negative solid tumors. Nat Immunol 24, 1345–1357 (2023). 10.1038/s41590-023-01543-9

46 Chang, F. et al. Immune marker expression of irradiated mesothelioma cell lines. Front Oncol 12, 1020493 (2022). 10.3389/fonc.2022.1020493

47 Evans, R. A. et al. Lack of immunoediting in murine pancreatic cancer reversed with neoantigen. JCI Insight 1, pii: e88328 (2016). 10.1172/jci.insight.88328

48 Melero, I., Castanon, E., Alvarez, M., Champiat, S. & Marabelle, A. Intratumoural administration and tumour tissue targeting of cancer immunotherapies. Nat Rev Clin Oncol 18, 558–576 (2021). 10.1038/s41571-021-00507-y

49 Hundal, J. et al. pVAC-Seq: A genome-guided in silico approach to identifying tumor neoantigens. Genome Med 8, 11 (2016). 10.1186/s13073-016-0264-5

50 Zheng, G. X. et al. Massively parallel digital transcriptional profiling of single cells. Nat Commun 8, 14049 (2017). 10.1038/ncomms14049

51 Hao, Y. et al. Dictionary learning for integrative, multimodal and scalable single-cell analysis. Nat Biotechnol 42, 293–304 (2024). 10.1038/s41587-023-01767-y

52 Huang, L. C. et al. scDemultiplex: An iterative beta-binomial model-based method for accurate demultiplexing with hashtag oligos. Comput Struct Biotechnol J 21, 4044–4055 (2023). 10.1016/j.csbj.2023.08.013

53 Verstaen, K. et al. DALI (Diversity AnaLysis Interface): a novel tool for the integrated analysis of multimodal single cell RNAseq data and immune receptor profiling. bioRxiv, 2021.2012.2007.471549 (2022). 10.1101/2021.12.07.471549

54 Valkiers, S., Van Houcke, M., Laukens, K. & Meysman, P. ClusTCR: a python interface for rapid clustering of large sets of CDR3 sequences with unknown antigen specificity. Bioinformatics 37, 4865–4867 (2021). 10.1093/bioinformatics/btab446

55 McDavid, A., Finak, G. & Yajima, M. MAST: Model-based Analysis of Single Cell Transcriptomics. (2025). doi:10.18129/B9.bioc.MAST

56 Zhou, Y. et al. Metascape provides a biologist-oriented resource for the analysis of systems-level datasets. Nat Commun 10, 1523 (2019). https://doi.org/10.1038/s41467-019-09234-6 [pii] 9234 [pii] 10.1038/s41467-019-09234-6

